# Extraction workflow determines marker-specific recovery and reproducibility in leaf-litter eDNA metabarcoding

**DOI:** 10.64898/2026.07.15.738510

**Authors:** Sven Weber, Pritam Banerjee, Edoardo Scali, Alex Farrow, Alex Boren, Tommy W. Russell, Natalie R. Graham, Rosemary G. Gillespie, George K. Roderick

## Abstract

Forest-floor leaf litter is a dynamic and structurally complex ecological transition zone and thus a promising substrate for terrestrial eDNA metabarcoding. Yet, extraction workflows for this heterogeneous matrix remain poorly standardized, especially in tropical systems, making it largely impossible to compare ecological functions across space, time, and taxa. To guide workflow selection across a series of selection criteria, including biological target, research question and practical considerations, we compared DNA extraction workflows for leaf-litter eDNA collected from 42 biological samples across seven different forest sites on Oʻahu, Hawaiʻi. We evaluated four DNA extraction workflows: (1) Two low-volume approaches, with DNA extracted directly from 200 mg of homogenized litter using (i) CTAB or (ii) DNeasy PowerSoil®; and (2) two high-volume approaches using PBS wash-based from 10 g of litter followed by (i) Centrifugation or (ii) Filtration. Taxonomic recovery from each workflow was evaluated with two COI primer sets targeting arthropods (ANML and shorter NoPlant), and one ITS marker targeting fungi. Results show that eDNA workflows tested here recovered site-level differences among forest-floor communities, but biodiversity recovery depended strongly on extraction workflow and marker. For low volumes, PowerSoil recovered the highest fungal richness (with ITS marker), and produced the most reproducible PCR-replicate profiles across markers, and required the least hands-on time, while CTAB was less expensive but required handling hazardous chemicals. For high volumes workflow, Centrifugation recovered higher arthropod diversity with ANML primer. Differences in community composition were nonetheless recovered by each method. At the same time, sampling sites explained more ASV-level compositional variation than extraction workflow across markers, showing that all workflows retained site-level ecological signals. Together, these results support a workflow framework in which extraction choice depends on target organism group, DNA state, reproducibility needs, and practical constraints.

## 1. Introduction

Just above the soil surface, leaf litter supports diverse assemblages of fungi, bacteria, arthropods, and other metazoa that contribute to decomposition, nutrient cycling, soil formation, and the turnover of organic matter (Anderson, 1975; Osono, 2007) and represents the transition zone between living vegetation and mineral soil (Prescott & Vesterdal, 2021). Despite this ecological importance, leaf-litter biodiversity and its food web structure remain poorly understood, partly because these communities are physically difficult to sample and monitor using conventional sampling approaches. In addition, many litter-associated organisms are small-bodied, cryptic, seasonal, taxonomically unresolved, or present as immature life stages (Basset et al., 2020, 2022; Young & Hebert, 2022). For these reasons, leaf litter is a particularly promising matrix for terrestrial eDNA surveys (Lopes et al., 2021; Yang et al., 2014), especially in tropical ecosystems, where biodiversity is high, taxonomic impediments are substantial, and tools for rapid assessment are increasingly needed for conservation, invasive-species management, and restoration.

Despite the promise, leaf litter is a challenging substrate for eDNA. Unlike water, where DNA is suspended in a relatively homogeneous medium, or soil, where extraction protocols have been more extensively tested and optimized, leaf litter is physically and biologically heterogeneous at fine spatial scales (Lopes et al., 2021; Mauvisseau et al., 2022). DNA in this matrix is unlikely to occur in a single state: it may be intracellular, extracellular, particle-bound, protected inside spores, associated with frass, or present in degraded tissue fragments (Hermans et al., 2022; Zinger et al., 2016). These fractions may differ in persistence, accessibility, inhibitor exposure, and amplification success. As a result, leaf-litter eDNA studies to date have used extraction workflows that vary substantially in fundamental aspects, including, input mass, methods of homogenization, extraction chemistry, inhibitor removal, and whether DNA is recovered directly from litter or indirectly from wash solutions (Barnes & Turner, 2016; Basset et al., 2020; Castillo et al., 2026). Low-volume direct extraction approaches, such as CTAB-based protocols or commercial soil kits like the PowerSoil® DNA Isolation kit, are relatively standardized and scalable, but they process only a small aliquot of a heterogeneous substrate (100-200mg). High-volume wash-based methods can integrate more material and reduce subsampling effects, but they also require more handling time, equipment, reagents, and may introduce DNA loss and contamination. Coupled with the volume, the taxonomic diversity recovered from leaf litter depends on the DNA primers used (Elbrecht & Leese, 2017; Krehenwinkel, Weber, Künzel, et al., 2022) because these differ in taxonomic target, amplification bias, and reference-database structure (Deagle et al., 2014; Schoch et al., 2012; Tedersoo et al., 2022).

To simplify the choice of options for leaflitter eDNA and to match possible workflows for leaflitter eDNA with project goals and feasibility, we compared four DNA extraction workflows for leaf-litter eDNA metabarcoding across seven forest sites on Oʻahu using two broad strategies and two methods within each: (1) low-volume direct extraction from homogenized litter with (i) CTAB and (ii) DNeasy PowerSoil; and (2) high-volume wash-based recovery with (i) centrifugation and (ii) filtration. Within each method we evaluated three primer sets targeting different taxa – two COI markers for arthropods and one ITS marker for fungi – assessing taxonomic recovery, reproducibility, and practical feasibility. We conducted four linked analyses. **First**, because low- and high-volume workflows process different amounts of litter and recover DNA through different mechanisms, we tested whether workflows differed in read yield, sample retention after filtering and rarefaction, and taxonomic recovery. **Second**, we tested whether these differences were marker and thus taxon-dependent, expecting high-volume workflows to favor spatially patchy arthropod DNA more than fungal DNA. **Third**, we asked whether extraction workflow altered ecological inference by comparing workflow effects with among-site community differences and by quantifying within-sample PCR-replicate reproducibility. **Fourth**, we compared cost, workload, and cost per successful sample among methods. Together, these analyses evaluate extraction workflows for leaf litter eDNA as biological and practical trade-offs rather than as a search for a single universally best method. We synthesize these results into a practical decision framework for selecting leaf-litter eDNA extraction workflows (**Fig. 6)**.

## 2. Material and Methods

### 2.1. Field sampling

Leaf-litter sampling was conducted in July 2023 and January 2024 at seven forest sites across Oʻahu, Hawaiʻi (**Fig. 1**). Sites were located in four geographic contexts: Kolekole, represented by Puʻu Hapapa and Kaluaʻa Ridge; the Kahuku region, represented by Kahuku Trail and Drum Road; the Waikāne region, represented by Schofield-Waikāne and Centerline Road; and Haleʻauʻau Gulch. The sites spanned a gradient from higher-elevation, relatively intact forests with greater native tree cover to lower-elevation, more degraded forests with higher representation of invasive vegetation, with Haleʻauʻau Gulch representing a mixed forest.

**Figure 1.**
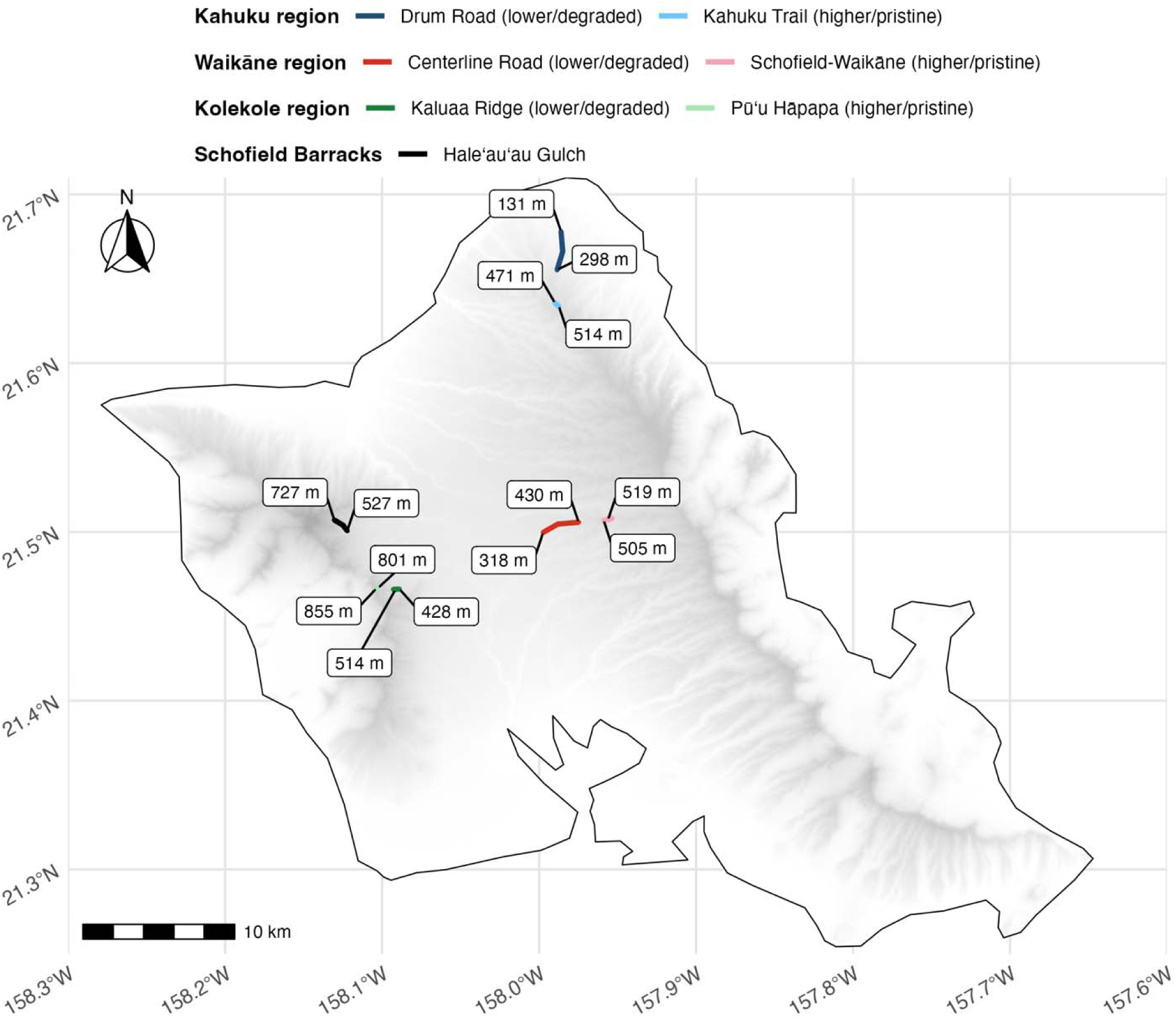
Study design and spatial layout of leaf-litter sampling sites on Oʻahu. Sampling sites were grouped into four geographic contexts: the Kahuku region, Waikāne region, Kolekole region, and Haleʻauʻau Gulch. Colored line segments indicate site locations and pairings used in the extraction-method comparison, with lower/degraded and higher/pristine sites shown within regions where applicable. Elevation labels denote the sampled range within each site

At each site, three sampling locations were established, and two biological field replicates were collected from adjacent 1 m² quadrats at each location, yielding 42 biological samples in total. We collected surface leaf litter down to the soil boundary. Approximately 500 g of litter from each quadrant was coarsely sieved using a litter reducer with 10 mm mesh (EntomoAlex-GR, Italy) to remove large debris such as twigs and branches. Samples were packed into commercially available ziplock bags, transported to the field station on Oʻahu immediately after collection, and processed for preservation before shipment. Leaf-litter samples were freeze-dried for at least 48 h, or until all visible moisture had been removed, using a Labconco FreeZone® 1 Liter Benchtop freeze dryer (Model 7740020, Labconco, USA). Particularly damp samples required extended drying times of up to four days. Freeze-drying was used to stabilize the material and reduce DNA degradation during subsequent handling and transport (Weber et al., 2026).

### 2.2 DNA extraction workflows

We compared four extraction workflows grouped into two broad strategies. (1) Low-volume workflows extracted DNA directly from 200 mg of homogenized leaf litter using either (i) a CTAB-based protocol or (ii) the DNeasy PowerSoil® kit. (2) High-volume workflows processed 10 g of intact leaf litter through a PBS wash, followed by DNA recovery from the wash solution using either (i) centrifugation or (ii) filtration.

### 2.3 Low-volume extractions

For the low-volume extraction methods, dried leaf litter was homogenized in a household blender (Ninja®), following previous plant and litter-associated eDNA workflows (Weber et al., 2024, 2026). To minimize cross-sample contamination, the blender blades and container were cleaned after each sample using a three-step cleaning protocol adapted from (Buchner et al., 2021). Homogenized material was transferred into one or two sterile 50 ml Falcon tubes, depending on sample volume, and stored at -20°C until DNA extraction (Stothut et al., 2024). The same homogenized material was used as source material for both low-volume workflows: CTAB (OPS Diagnostics LLC, New Jersey, USA) and DNeasy PowerSoil (Qiagen, California, USA) for low-volume extraction methods. For the CTAB extraction we followed the standard CTAB protocol (Doyle & Doyle, 1987), with slight modifications after Krehenwinkel et al. (2022). 200 mg of homogenized leaf litter was transferred to a 2 ml polypropylene screw-cap tube containing two sterile 2.3 mm metal beads. Samples were bead-beaten for 30 s twice on a SPEX bead beater (Spex SamplePrep MiniG Tissue Homogenizer, New Jersey, USA) at 1100 rpm. Subsequently, 1 mL CTAB extraction buffer was added and samples were incubated at 60°C for 30 min in an incubator (Yamato Scientific, California, USA). After incubation, samples were centrifuged for 5 min at 14,000 × g, and the supernatant was transferred to a fresh 1.5 mL reaction tube. This clarification step was repeated once to further reduce carryover of fine particulates. An equal volume of 25:24:1 phenol:chloroform:isoamyl alcohol (Sigma Aldrich, Massachusetts, USA) was then added, followed by centrifugation for 1 min at 14,000 × g. The upper aqueous phase was transferred to a new 1.5 mL reaction tube, mixed with 0.7 volumes of chilled 100% isopropanol, and incubated at −20°C for 15 min to precipitate DNA. Samples were then centrifuged for 10 min at 14,000 × g, the supernatant was discarded, and the pellet was rinsed with 70% ethanol, dried in a SpeedVac (Eppendorf, Hamburg, Germany), and resuspended in 35 µL hydration solution (Qiagen, California, USA). For the DNeasy PowerSoil workflow, the manufacturer’s protocol was followed with minor modifications. Briefly, 200 mg of homogenized leaf litter was used as input. The initial vortexing step was replaced by two bead-beating rounds on the SPEX bead beater (Spex SamplePrep MiniG Tissue Homogenizer, New Jersey, USA), and DNA was eluted in 50 µL of pre-heated Solution C6 after a 10 min incubation prior to centrifugation.

### 2.4 High-volume extractions

For the high-volume workflows, 10 g of intact, non-homogenized leaf litter was incubated in a 10 L Pyrex bottle containing 1.5 L of 1× PBS (Thermo Fisher Scientific, Massachusetts, USA) and 3 mL of 1 N HCl. Samples were incubated on a heated stir plate at 50°C for approximately 18 h and then left undisturbed overnight at room temperature to allow larger particles to settle. The supernatant was passed through a coffee filter to remove coarse debris, and the resulting filtrate served as the shared source material for both the centrifugation and filtration workflows.

For the centrifugation workflow, four 50 mL aliquots of filtrate from the same wash source were centrifuged for 10 min at 14,500 × g (Sorvall X4R Pro-MD Benchtop Centrifuge, Thermo Fisher Scientific, Massachusetts, USA). After centrifugation, the supernatant was discarded and the remaining pellet was dried at 56°C overnight in an incubator (Yamato Scientific, California, USA). DNA was extracted from the dried pellet using a modified DNeasy Blood & Tissue protocol after Banerjee et al. (2025), with the initial lysis step performed in a 50 mL Falcon tube. Although both high-volume workflows originated from the same 10 g litter wash, the effective extracted filtrate fraction differed between workflows. Filtration processed 300 mL total filtrate across three filters, but only one half of each filter was extracted, corresponding to approximately 150 mL equivalent filtrate. Centrifugation extracted pellets generated from 200 mL total filtrate across four 50 mL tubes.

For the filtration workflow, three 100 mL aliquots of filtrate were passed through disposable 2.3 cm diameter, 0.45 µm pore-size filters (3 × 100 mL; Nalgene™ Single Use Analytical Filter Funnels, Thermo Fisher Scientific, Massachusetts, USA) using vacuum filtration for each sample. After filtration, each filter was divided into two halves: one half was used for DNA extraction and the other half was archived in Longmire’s buffer (Longmire et al., 1997). DNA extraction from filter material was performed using DNeasy Blood & Tissue kits (Qiagen, California, USA) with minor protocol modifications after Banerjee et al. (2025).

### 2.5 Contamination controls

Multiple controls were included to assess contamination during DNA extraction, PCR amplification, and laboratory handling. First, a DNA extraction blank was included for each extraction workflow and processed alongside samples (Salter et al., 2014; Tedersoo et al., 2022). Second, negative PCR controls were included during amplification for all primer sets. Third, we conducted a dedicated laboratory airborne-exposure experiment to evaluate background fungal contamination in the Berkeley laboratory environment. Because ITS primers can amplify ubiquitous fungal DNA from laboratory air, surfaces, and human-associated sources, open tubes containing approximately 1.5 mL DNA-grade water were exposed to laboratory air for different durations before being closed. Exposure times were 0 min, 1 min, 5 min, 15 min, 30 min, 1 h, 2 h, 3 h, 4 h, 5 h, 6 h, 1 day, 2 days, 3 days, and 10 days. These exposed water controls were carried through the ITS amplification workflow to assess whether the laboratory environment was suitable for fungal metabarcoding work.

### 2.6 PCR amplification and library preparation

Extracted DNA was initially assessed by NanoDrop (Thermo Fisher Scientific, Massachusetts, USA) & qubit 3.0 fluorometer (Thermo Fisher Scientific, USA) to estimate concentration and purity. Extracts exceeding 200 ng/µL were diluted with molecular-grade water prior to PCR to standardize template input and reduce the risk of PCR inhibition or nonspecific amplification associated with excessive template DNA and co-extracted inhibitors (McKee et al., 2015; Sidstedt et al., 2020). The analysis included 42 biological samples, four extraction workflows (CTAB, DNeasy PowerSoil, filtration, and centrifugation), and three marker assays (ANML, NoPlant, and ITS), yielding 504 sample-by-workflow-by-marker combinations. Each combination was amplified in three PCR replicates, resulting in 1,512 PCR reactions before replicate pooling.

All PCRs were performed in 10 µL reactions containing 5 µL 2× QIAGEN Multiplex PCR Master Mix (Qiagen, Germany), 1 µL DNA template, 0.5 µL of each primer (10 µM), 1 µL Q-solution, and 2 µL nuclease-free water. Thermal cycling consisted of an initial denaturation at 95°C for 15 min, followed by 35 cycles of 94°C for 30 s, 46°C for 90 s, and 72°C for 90 s, with a final extension at 72°C for 10 min. Amplification success and expected fragment length were checked by 2% agarose gel electrophoresis with GelRed (Fisher Scientific, USA), and technical PCR replicates were subsequently pooled based on relative gel-band intensity. Indexed libraries were generated in a second PCR step to attach dual 8 bp indices and Illumina adapter sequences (Lange et al., 2014). Sample pooling was based on relative band intensity on 2% agarose gels stained with GelRed (Fisher Scientific, USA). Amplicons were pooled by primer set, resulting in one sequencing pool per primer. Each pool was purified using standard bead selection with 0.8× volume of AMPure XP beads (Beckman Coulter, USA). Library pools were quantified using an Invitrogen Qubit 3.0 fluorometer (Thermo Fisher Scientific, USA), and fragment size was assessed on an Agilent Bioanalyzer 2100 system (Agilent Technologies, USA) at QB3 Genomics, University of California, Berkeley. Primer-specific pools were then quantified by qPCR at the pool level and normalized by molarity to generate the final sequencing superpool, targeting 150,000 reads per sample across the ANML, NoPlant, and ITS assays. Sequencing was performed at QB3 Genomics, University of California, Berkeley, on a single NextSeq 2000 run using P2 600-cycle paired-end chemistry with a 30% PhiX spike-in.

### 2.7 Bioinformatic processing

Raw sequence data were demultiplexed using BCL Convert with zero mismatches allowed in the dual-index combinations. All paired-end reads were merged using PEAR with a minimum overlap of 50 bp and a minimum quality score of 20 (Zhang et al., 2014). Reads were further filtered to retain sequences with Q ≥ 33 across more than 90% of positions and converted to FASTA format. Primer sequences and inline barcodes were removed using a custom Unix-based pipeline allowing for degenerate bases (Weber et al., 2026). Exact amplicon sequence variants were inferred using VSEARCH with a minimum cluster size of eight and de novo chimera removal. We refer to these sequence units as ASVs throughout, recognizing that they represent error-corrected amplicon sequences rather than taxonomic species (Rognes et al., 2016). PCR replicate concordance was evaluated after sequencing. Replicate sets were merged when they shared at least an R² of 0.75 (Pearson correlation), and samples were retained only when at least two of three replicate libraries were successful. Libraries were subsequently rarefied to 3,000 reads per sample for downstream comparisons, while unrarefied data were retained for reference (**Fig. S1 and Fig. S2**) (Schloss, 2024). For fungal data, ASVs were queried against the UNITE database updated in September 2025 for taxonomic assignment (Abarenkov et al., 2024). For arthropod data (ANML and NoPlant), ASVs were queried against the NCBI nt database downloaded in February 2025 using BLASTn, retaining a maximum of 10 hits per query (Camacho et al., 2009). Taxonomic information was parsed using a custom Python workflow based on blast2tax (Schöneberg, 2023). Non-target reads (non-arthropod identifications) were removed, and only arthropod ASVs were retained for downstream analyses of COI data. For ANML and NoPlant, taxonomic filtration was done on the following criteria to eliminate all potential contamination: Sequences were required to have a minimum length of 170 bp for the 180-bp ANML amplicon and 64 bp for the 74-bp NoPlant amplicon, corresponding to a ≥90% length match against the Blast reference.

All downstream data handling, filtering, visualization, and statistical analyses were conducted in R 4.5.1 (R Core Team, 2024). Analyses were run separately for ANML, NoPlant, and ITS. Libraries retained for downstream analysis were rarefied to 3,000 reads per sample-by-workflow-by-marker library before diversity comparisons. We calculated rarefied observed ASV richness and Shannon diversity for each retained library and compared extraction workflows using Kruskal-Wallis tests followed by Benjamini-Hochberg-adjusted Dunn post hoc tests (Benjamini & Hochberg, 1995). Read yield was analyzed separately from rarefied diversity metrics because it was measured before rarefaction. Because each biological leaf-litter sample was processed with each of the four extraction workflows, read yield was treated as a paired response across extraction workflows. Friedman tests were used for paired read-yield and PCR-replicate reproducibility summaries. Tests were conducted only on complete biological blocks, defined as biological samples represented by all four extraction workflows for the relevant marker and response; incomplete blocks were excluded from the corresponding test, and no values were imputed. In the current read-yield and reproducibility Friedman tests, all three markers retained 42 complete blocks. As secondary sensitivity analyses, we fitted Gaussian linear mixed-effects models with extraction workflow as a fixed effect and biological sample as a random intercept. For read yield, the response was log1p-transformed post-filtering read count; for rarefied observed richness, the response was log1p-transformed richness; and for Shannon diversity the response was untransformed Shannon diversity. These Gaussian linear mixed-effects models were fitted with lme4::lmer() using random intercepts for biological sample and for the biological-sample-by-workflow unit, and workflow effects were tested by likelihood-ratio chi-square tests comparing models with and without extraction workflow. For sequencing-success analyses, a library was defined as a sample-by-workflow-by-marker unit after demultiplexing, marker-specific filtering, and pooling of the three PCR replicates. Sequencing success was evaluated at ≥1,500 and ≥3,000 post-filtering reads per pooled library; the 3,000-read threshold matched the rarefaction depth used for diversity and community-composition analyses. PCR-replicate-level rarefaction curves were inspected separately with a 1,500-read reference line, but the success models used pooled library totals rather than individual PCR-replicate totals. Sequencing success was analyzed with binomial GLMMs in lme4::glmer() ; marker-specific success models included extraction workflow as a fixed effect and random intercepts for biological sample and biological-sample-by-workflow unit, with workflow effects evaluated by likelihood-ratio tests and pairwise workflow contrasts estimated with emmeans() using Benjamini-Hochberg correction. Community composition was assessed with PERMANOVA using site and extraction workflow as predictors, with betadisper dispersion checks, pairwise PERMANOVA contrasts, and NMDS stress values reported for ordination quality. Taxonomic-composition comparisons used only balanced biological units represented across all four workflows, applied a ≥90% identity filter, and tested paired presence and relative-abundance differences with Cochran’s Q, exact McNemar, and paired Wilcoxon tests with Benjamini-Hochberg correction. Cost and efficiency summaries were calculated per extraction workflow, with bootstrap confidence intervals for cost per successful sample and cost per richness unit (Bates et al., 2015; Oksanen et al., 2013).

## 3. Results

### 3.1 Sequencing output across extraction methods and markers

After quality control, sample retention differed among extraction workflows and markers (**Table 3; Fig. S1–S2**). PowerSoil and filtration retained the highest number of samples across markers, followed, by CTAB and centrifugation, across all the markers (**Table 3**). Read yield showed marker-dependent workflow effects (**Table 3; Supplement Table S2**). ANML read yield did not differ among workflows, NoPlant showed the clearest workflow effect, and ITS showed a workflow effect in the paired Friedman test but not in the mixed-model sensitivity analysis. PCR-replicate reproducibility differed among extraction workflows across all markers (**Fig. S8; Table 3; Supplement Table S2**), with PowerSoil showing the most reproducible PCR-replicate profiles compared with CTAB, centrifugation, and filtration. Library-preparation exposure controls showed increasing read counts with longer exposure duration, whereas observed fungal richness in controls remained low across exposure times (**Fig. S3**).

**Table 1.** Overview of DNA extraction workflows tested for leaf-litter eDNA.

| Method | Extraction strategy | Input mass (g) | Material state | Recovery principle | Extraction chemistry / kit | Cost per sample (USD) | Hands-on time per batch (h) |
| --- | --- | --- | --- | --- | --- | --- | --- |
| CTAB | low-volume direct | 0.2 | homogenized litter | direct lysis | CTAB chemistry | 3.15 | 2.62 |
| PowerSoil | low-volume direct | 0.2 | homogenized litter | bead beating | DNeasy PowerSoil kit | 10.55 | 1.75 |
| Centrifugation | high-volume wash-based | 10 | intact litter/wash | buffer wash plus centrifugation | Wash, centrifugation, extraction kit | 22.64 | 2.19 |
| Filter | high-volume wash-based | 10 | intact litter/wash | buffer wash plus filtration | Wash, filtration, extraction kit | 35.16 | 3.06 |

**Table 2.**
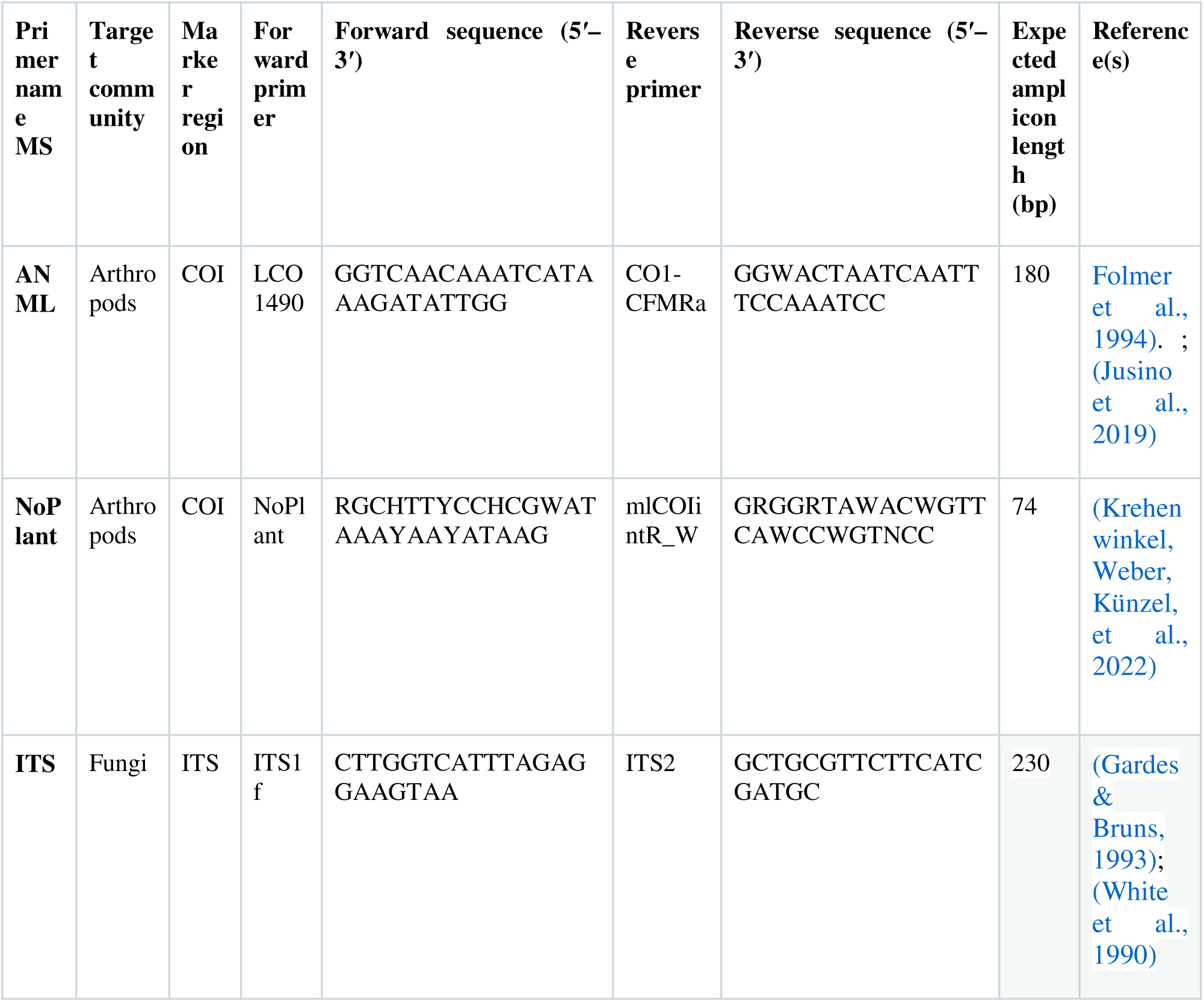
Primer-pairs used in the study.

**Table 3.** Sequencing performance, diversity, reproducibility across markers and extraction methods.

| Marker | Method<br>(n =42) | Retained<br>(n) | Below<br>rarefaction<br>% | Mean<br>Reads,<br>[IQR] | Mean<br>observed<br>richness,<br>[IQR] | Shannon<br>diversity,<br>[IQR] | Median replicate<br>dissimilarity |
| --- | --- | --- | --- | --- | --- | --- | --- |
| ANML | CTAB | 35 | 16.7 | 11600 [4178-21306] | 109.0<br>[76.0-144.5] | 2.42 [1.88-2.97] | 0.382 |
| ANML | PowerSoil | 41 | 2.4 | 16215 [6926-29830] | 92.0 [75.0-119.0] | 2.43 [2.10-2.65] | 0.202 |
| ANML | Centrifugation | 32 | 23.8 | 12268 [3488-18206] | 144.0<br>[132.8-173.2] | 2.83 [2.45-3.35] | 0.337 |
| ANML | Filter | 39 | 7.1 | 14448 [9724-23464] | 81.0 [64.5-96.5] | 2.56 [2.05-2.89] | 0.348 |
| NoPlant | CTAB | 29 | 31.0 | 9420 [2702-17318] | 85.0 [64.0-111.0] | 2.56 [2.26-2.97] | 0.383 |
| NoPlant | PowerSoil | 41 | 2.4 | 17765 [8796-35342] | 84.0 [67.0-119.0] | 2.40 [1.95-2.81] | 0.216 |
| NoPlant | Centrifugation | 32 | 23.8 | 6220 [3334-13986] | 83.5 [66.0-99.0] | 2.19 [1.78-2.62] | 0.366 |
| NoPlant | Filter | 41 | 2.4 | 14069 [8810-24899] | 76.0 [53.0-100.0] | 2.31 [1.86-2.78] | 0.310 |
| ITS | CTAB | 33 | 21.4 | 37780 [13658-47816] | 349.0<br>[279.0-425.0] | 4.23 [3.75-4.66] | 0.284 |
| ITS | PowerSoil | 42 | 0.0 | 50948 [45006-59474] | 437.5<br>[407.0-481.2] | 4.62 [4.47-4.87] | 0.199 |
| ITS | Centrifugation | 30 | 28.6 | 16806 [1572-31180] | 337.5<br>[222.8-437.2] | 3.90 [3.57-4.68] | 0.360 |
| ITS | Filter | 41 | 2.4 | 34483 [28014-40233] | 335.0<br>[252.0-393.0] | 4.32 [3.88-4.69] | 0.332 |

### 3.2 ASV richness differs among extraction methods

After quality control, ASV richness differed among extraction workflows, and markers (**Fig. 2**; **Table 3; Supplement Table S2**). Workflow effects were strongest for ANML (Kruskal-Wallis χ² = 47.36, df = 3, p < 0.001; mixed model χ² = 42.03, p < 0.001) and ITS ((Kruskal-Wallis χ² = 26.56, df = 3, p < 0.001; mixed model χ² = 75.75, p < 0.001), whereas NoPlant showed weaker richness differences (**Table 3; Figure S5**). In case of ANML primer, centrifugation recovered the highest ASV richness and unique ASVs followed by CTAB, PowerSoil and filtration (Table 3; Figure S5). For ITS, PowerSoil detected the highest ASVs richness and unique ASVs, followed by CTAB, Filtration and Centrifugation (**Table 3; Fig. S5**). For the NoPlant primer richness differences among workflows were weaker, PowerSoil and Filtration detected the highest unique ASVs compared to CTAB and Centrifugation (Figure S5).

**Figure 2.**
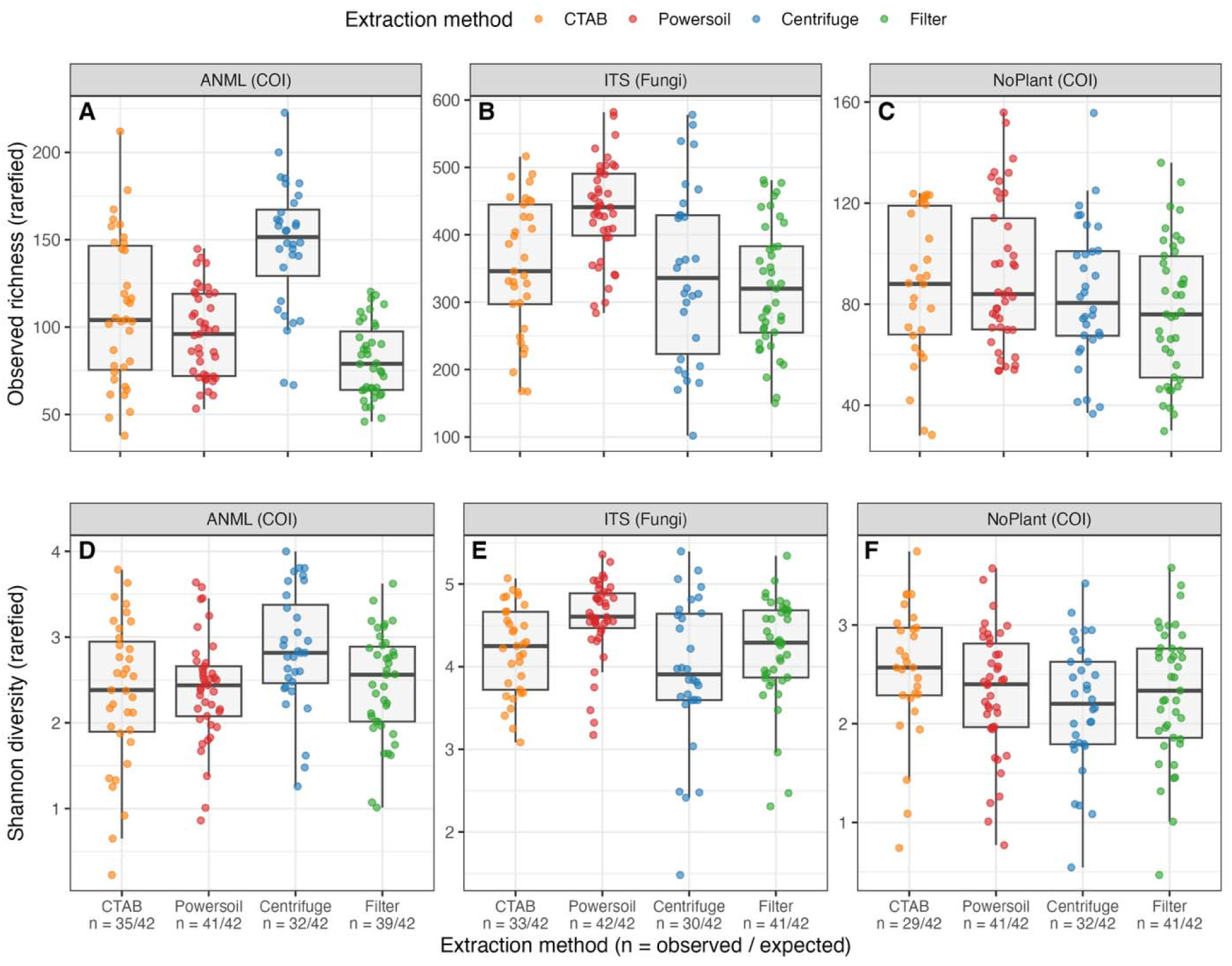
Rarefied alpha diversity based on observed richness/ASVs. **(A-C) and Shannon diversity (D-E) across extraction methods for three primer sets.** Rarefied richness differed among extraction methods for ANML (Kruskal-Wallis: χ² = 47.36, df = 3, p = 6.81 × 10^-10) and ITS (χ² = 26.56, df = 3, p = 2.10 × 10^-5), but not for NoPlant (χ² = 3.94, df = 3, p = 0.217).

### 3.3 Taxonomic and community composition vary by extraction workflow

Taxonomic composition showed variation across the workflow for each primer (**Fig. 3A; Fig. S6**). In case of ANML primer, Centrifugation showed largest number of pairwise differences, and detected several arthropod orders more frequently than CTAB, Filtration, or PowerSoil, including Lepidoptera, Amphipoda, Diptera, Collembola, and Hemiptera. For NoPlant, taxonomic differences among workflows were weaker than for ANML and ITS, with fewer order-level differences detected across extraction workflows (**Fig. 3B; Fig. S6**). In the case of ITS primer, PowerSoil showed the strongest taxonomic differences. Presence-absence recovery differed among workflows for 17 of 41 fungal classes after Benjamini-Hochberg correction, with major shifted fungal classes included Agaricomycetes, Dothideomycetes, Sordariomycetes, Eurotiomycetes, Mucoromycetes, and Mortierellomycetes.

**Figure 3.**
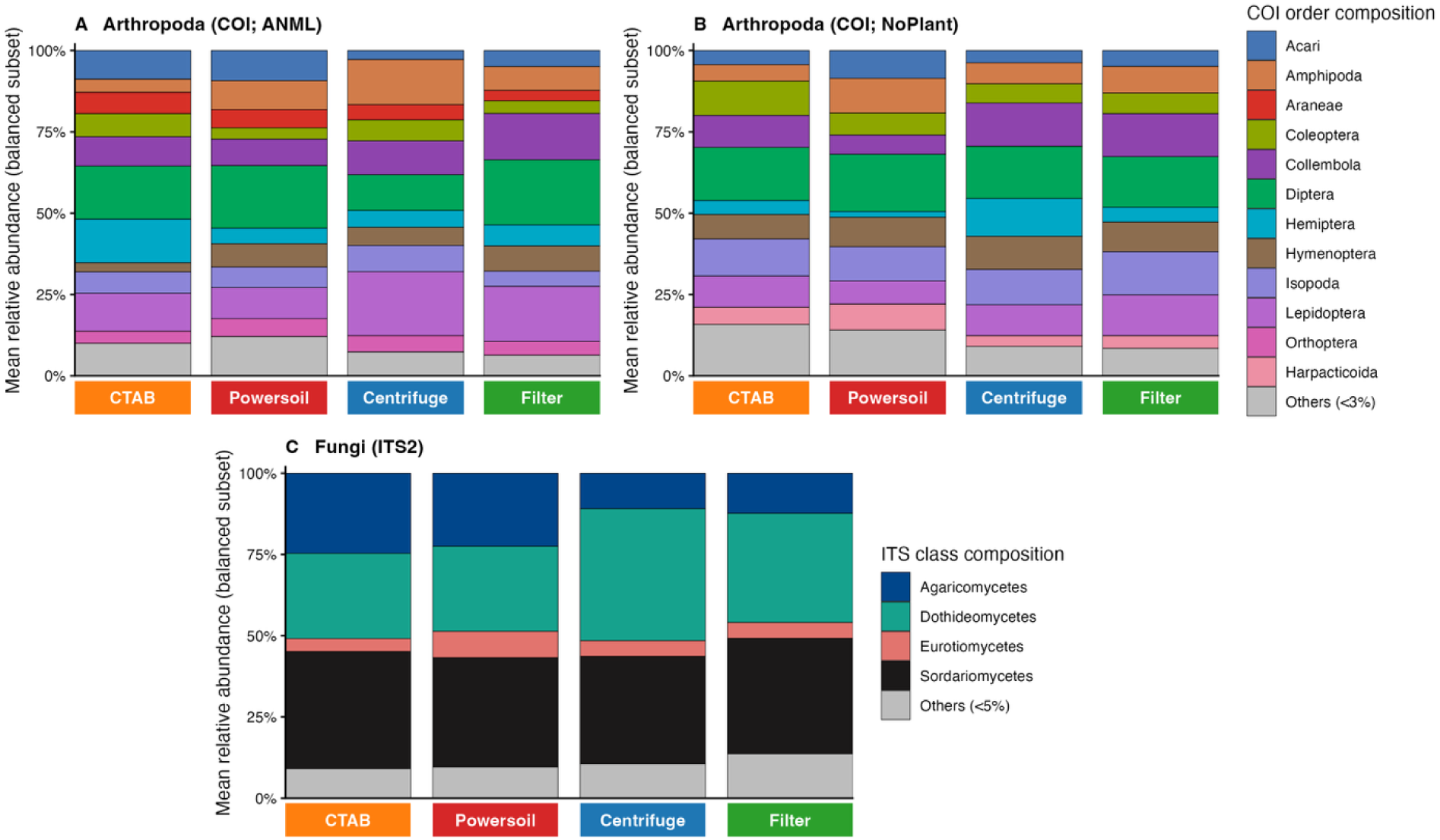
Taxonomic composition of the balanced subset across extraction methods. Stacked bars show mean relative abundance for the subset of biological sampling units represented across all four extraction methods, allowing direct method comparison without imbalance in sample representation. Arthropod composition is summarized at the order level for ANML and NoPlant, and fungal composition is summarized at the class level for ITS. Only dominant groups are shown explicitly; low-abundance taxa were pooled into an “Others” category.

Overall, community composition differed among extraction workflows across all three markers (**Fig. 4; Supplement Table S2**). PERMANOVA models detected significant workflow effects for ANML, NoPlant, and ITS across Hellinger-Bray, Jaccard, and Sørensen distances (**Supplemental Table 2)**. Workflow effects were largest for ANML, where extraction workflow explained 7.2–9.9% of compositional variation, compared with 2.6–3.1% for NoPlant and 2.6–3.3% for ITS. However, the sampling site explained more compositional variation than extraction workflow for every marker and distance metric (Fig. 4C). Dispersion analyses showed that workflow effects also involved differences in within-workflow variability (**Fig. S7; Supplement Table S2**). Under the paired permutation design, with permutations blocked by biological unit, betadisper tests detected significant dispersion differences among extraction workflows for ANML, NoPlant, and ITS across all three distance metrics.

**Figure 4.**
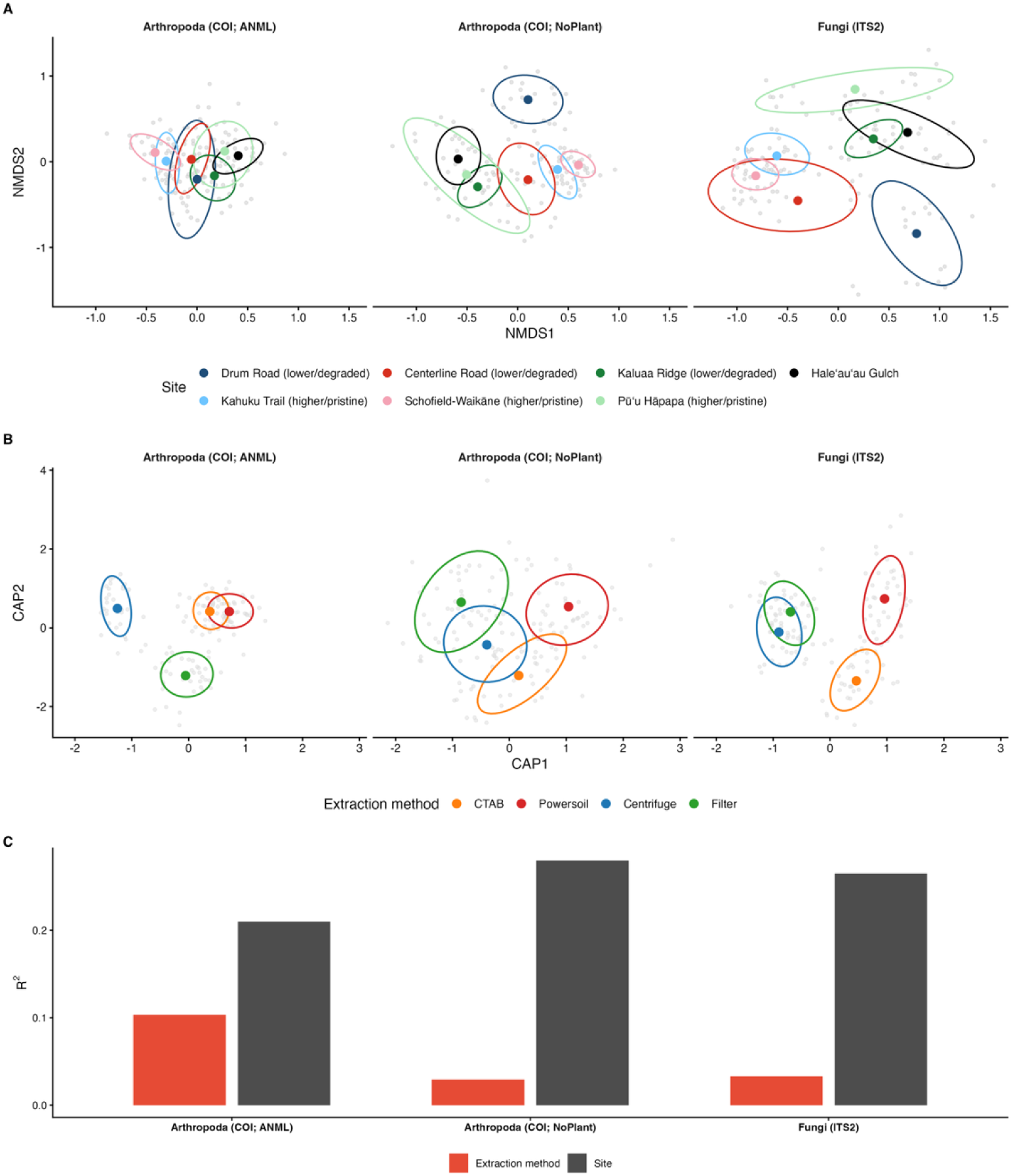
Community composition across sites and extraction methods. (A) Sørensen-based NMDS ordinations summarize site-level compositional structure for ANML, NoPlant, and ITS; colored centroids and ellipses denote site groups. (B) Partial CAP ordinations show extraction-method structure after conditioning on site, with colored centroids and ellipses denoting extraction methods. (C) Marginal PERMANOVA effect sizes from models including both site and extraction method. Across all three primer sets, the site explained more compositional variation than extraction method, although extraction method remained detectable. Marginal PERMANOVA models yielded smaller extraction-method effects (R² = 0.025 to 0.103) than site effects (R² = 0.201 to 0.279), with both terms significant in the fitted models (p = 0.001). These results indicate that the extraction method influenced recovery, but the site remained the stronger source of compositional variation.

### 3.4 Workflow trade-offs differ across recovery, reproducibility, cost, and practicalities

Extraction workflows differed across the decision axes summarized in the radar plots, including reproducibility, assay success, cost per sample, safety, yield, time per batch, and scalability potential (**Fig. 5**). Among low-volume methods, CTAB had the strongest cost profile ($3.15/sample), but lower safety scores because of phenol handling. PowerSoil was more expensive ($10.55/sample), but showed the broadest overall performance profile across markers, especially through high reproducibility, high assay success, and short hands-on time. Among high-volume methods, centrifugation showed a marker-specific ANML recovery profile, but at higher cost ($22.64/sample), whereas filtration showed high assay-success scores but weaker cost and scalability profiles. Overall, CTAB was best supported for cost-limited workflows (**Fig. 5A, E, I**), PowerSoil for reproducibility and processing efficiency, especially for ITS (**Fig. 5B, F, J**), centrifugation for ANML recovery (**Fig. 5C**), and filtration mainly for assay success and compatibility with filter-based workflows rather than for cost or overall recovery (**Fig. 5D, H, L**).

**Figure 5.**
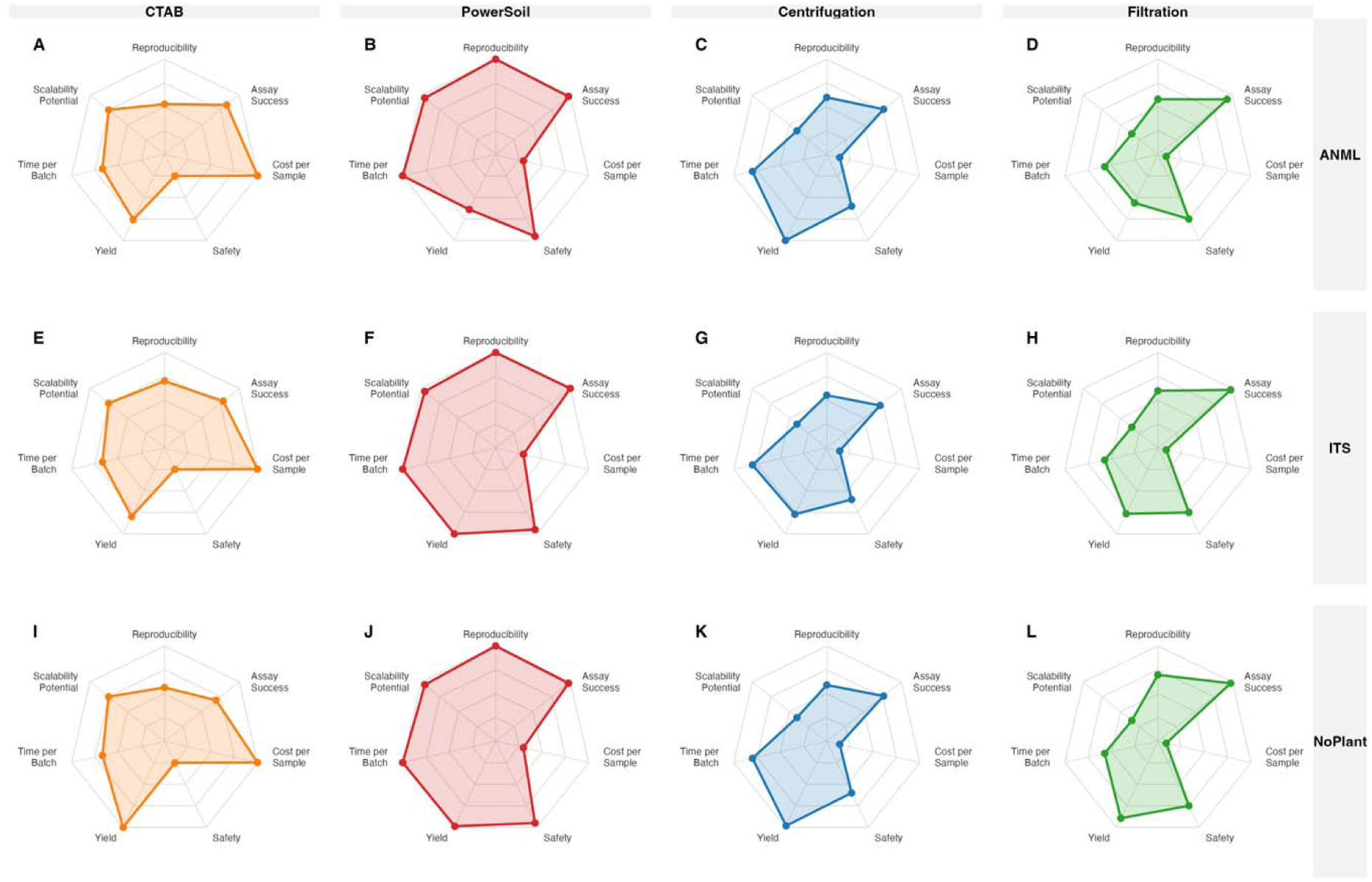
Decision-space comparison of leaf-litter eDNA extraction workflows across marker systems. Radar plots summarize the relative performance of four extraction workflows across seven decision axes: reproducibility, assay success, cost per sample, safety, yield, time per batch, and scalability potential. Columns represent extraction workflows: CTAB (A, E, I), PowerSoil (B, F, J), Centrifugation (C, G, K), and Filtration (D, H, L). Rows represent marker systems: ANML (A–D), ITS (E–H), and NoPlant (I–L). <u>All axes are oriented so that values farther from the center indicate more favorable performance</u>. Thus, larger filled areas indicate broader overall performance across the evaluated criteria. For practical axes, higher scores reflect more desirable conditions, such as lower cost, shorter processing time, or safer handling, rather than higher raw cost or longer processing time. Colors denote extraction workflows.

**Figure 6.**
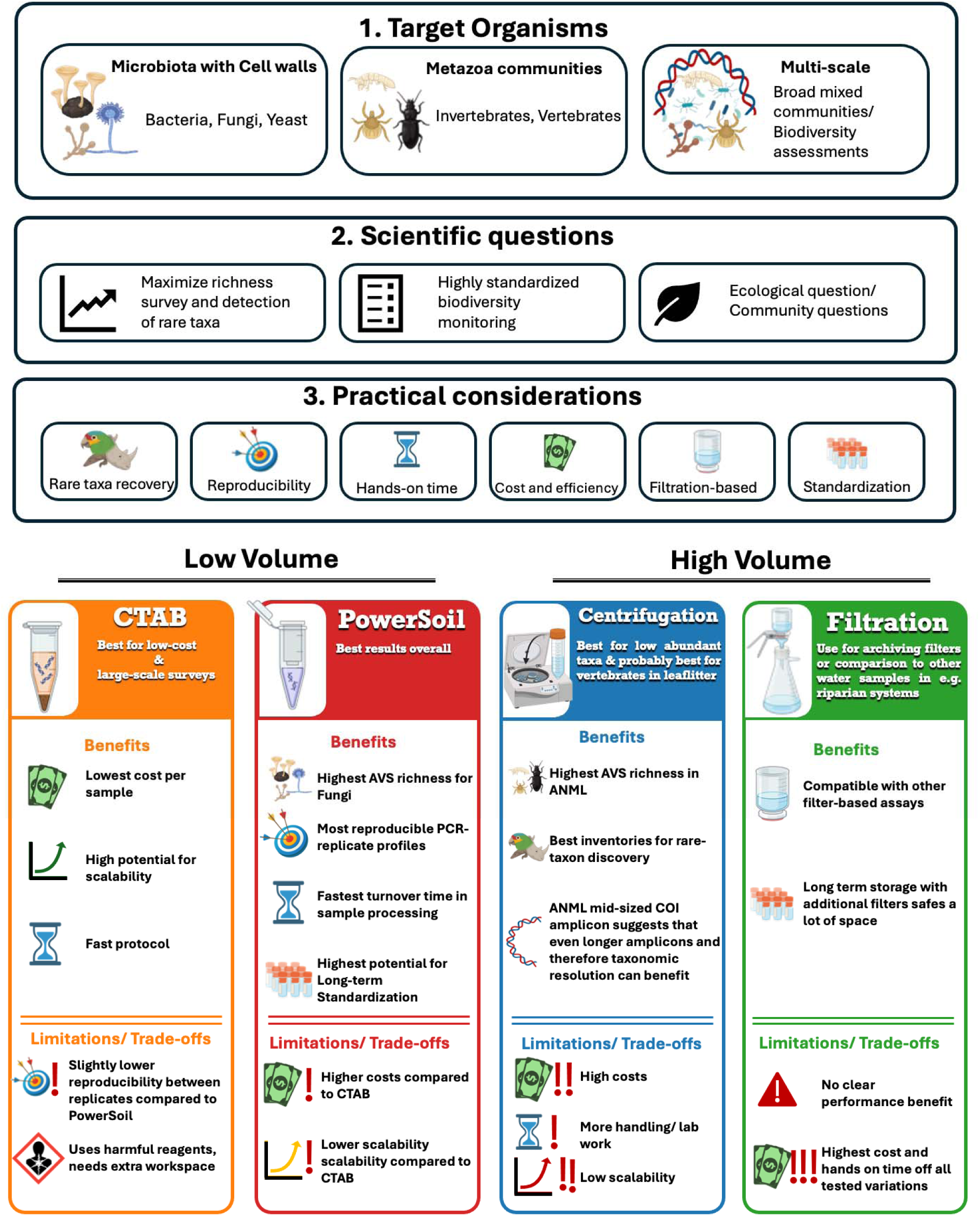
Practical decision framework for selecting leaf-litter eDNA extraction workflows. The flowchart summarizes how extraction-method choice can be guided by three sequential considerations: target organisms, scientific questions, and practical considerations. The upper section separates applications focused on microorganisms with cell walls, metazoan communities, or broader multi-taxon biodiversity assessment. The middle section distinguishes study objectives that prioritize richness and rare-taxon detection, standardized biodiversity monitoring, or community-level ecological comparisons. The lower section links these objectives to practical constraints, including reproducibility, processing time, cost, safety, scalability, filter compatibility, and long-term standardization. Method-specific panels summarize the principal advantages and limitations observed for (1) low volume (i) CTAB and (ii) PowerSoil; and (2) high volume (i) Centrifugation, and (ii) Filtration. For low volume, PowerSoil showed high assay success, reproducibility, and processing efficiency, with particularly strong performance for ITS-based fungal recovery. CTAB provides a lower-cost and scalable alternative but involves hazardous reagents and showed reduced reproducibility relative to PowerSoil. For high volume centrifugation showed the strongest recovery of ANML richness and is therefore most appropriate when rare arthropod or vertebrate detection is prioritized. Filtration is compatible with filter-based eDNA workflows and sample archiving but did not show a clear performance advantage in the present comparison. The framework is intended to support method selection according to study goals and constraints rather than to identify a universally optimal extraction workflow.

## 4. Discussion

In this study all leaf-litter eDNA workflows tested here recovered site-level differences among forest-floor communities, but biodiversity recovery depended strongly on extraction workflow and marker. No method was universally best across all criteria. Instead, workflow effects are best understood as a fraction-recovery problem: leaf litter contains extracellular, particle-associated, tissue-derived, and cellular DNA pools, and extraction workflows differ in which fractions they access most efficiently. High-volume centrifugation increased arthropod richness (with ANML primer) and order-level recovery, low-volume PowerSoil increased fungal richness (with ITS primer) and PCR-replicate reproducibility, and NoPlant was less affected by workflow than either ANML or ITS. At the same time, sampling sites explained more ASV-level compositional variation than extraction workflow across markers, showing that all workflows retained site-level ecological signals. Together, these results support a workflow framework in which extraction choice depends on target organism group, DNA state, reproducibility needs, and practical constraints.

### 4.1 Workflow affects sequencing success, and reproducibility

Leaf litter remains difficult to standardize as an eDNA substrate, due to its complex nature. As one of the main parts of eDNA detection is to accumulate the traces, the main problem with this kind of substrate is the low initial volume of samples for extraction (200 mg for low volume extraction). As we tested different volumes of extraction workflow, they will differ in input mass, lysis efficiency, inhibitor removal, and the recovery of DNA fraction (Barnes & Turner, 2016; Hermans et al., 2022; Mauvisseau et al., 2022; Zinger et al., 2016). This distinction matters because replicate variation is common in metabarcoding and eDNA studies, where low-template DNA, inhibitors, stochastic amplification, and primer-template mismatches can cause replicate libraries from the same sample to differ (Deagle et al., 2014; Elbrecht & Leese, 2017; Goldberg et al., 2016; Tedersoo et al., 2022). In our dataset, sequencing success, richness, and PCR-replicate reproducibility captured different aspects of workflow performance. PowerSoil produced the most consistent PCR-replicate profiles across all markers which, due to the standardized bead beating and inhibitor-removal chemistry, can improve reproducibility even when a workflow does not maximize richness. For leaf litter, inhibitor removal may be particularly relevant because decomposing plant material can contain humic substances, phenolics, and other PCR inhibitors (McKee et al., 2015; Sidstedt et al., 2020). However, processing of larger samples resulted in higher accumulation of animal DNA. Our comparisons provided reproducible results for each extraction method across different sampling locations.

### 4.2 Richness and marker-specific recovery

Although all workflow worked efficiently, high-volume extraction increased arthropod richness (with ANML primer) and low-volume PowerSoil increased fungal richness (with ITS primer) (Figure 2). This result is consistent previous work showing that terrestrial eDNA recovery depends on both extraction method and primer design, especially in heterogeneous substrates such as soil, litter, and dried plant material (Castillo et al., 2026; Horton et al., 2017; Krehenwinkel, Weber, Künzel, et al., 2022; Lopes et al., 2021; Ruppert et al., 2023). Among the high volume methods, Centrifugation worked better than Filtration method, although they used the same source of lysis solution (described in method). This may be due to higher chances of DNA accumulation in the Centrifugation workflow as it is avoiding the filter paper pore size parameter as we used in Filtration. Arthropod DNA in litter is likely sparse and unevenly distributed compared with microbial or fungal DNA, occurring as tissue fragments, frass, exuviae, extracellular DNA, or particle-associated DNA rather than dense cellular biomass. Thus, processing 10 g of litter through a wash-based workflow may therefore increase the chance of recovering diffuse animal traces relative to direct extraction from a 200 mg homogenized aliquot. This interpretation is consistent with broader eDNA work showing that DNA state, particle association, and sample-processing strategy influence detection (Barnes & Turner, 2016; Hermans et al., 2022; Mauvisseau et al., 2022; Zinger et al., 2016).

The effective detection arthropod with ANML primers also clarifies earlier concerns about COI in soil and leaf-litter eDNA. Horton, Kershner, and Blackwood (2017) found that COI recovered few invertebrate sequences from soil and leaf litter and thus they favored 18S for broad metazoan detection. Our study did not compare COI and 18S directly, so it cannot determine which marker is generally superior. However, our results show that ANML primer (COI) can recover high arthropod richness when paired with Centrifugation, suggesting that COI performance depends on primer design, amplicon length, sequencing depth, taxonomic filtering, and extraction workflow. This distinction is important because COI remains valuable for arthropod metabarcoding due to its taxonomic resolution and reference-database coverage, despite known primer-bias limitations (Deagle et al., 2014; Elbrecht & Leese, 2017).

On the other hand, low volume extraction, especially PowerSoil worked better for microbores, as a small portion of homogenized litter sample can contain millions of microbes, however higher volume should come with higher richness. Fungal DNA is often embedded in spores, hyphae, and decomposing plant tissue, and fungal metabarcoding is sensitive to extraction chemistry, mechanical disruption, inhibitor removal, where PowerSoil worked better than other methods (Lopes et al., 2021; Schoch et al., 2012; Tedersoo et al., 2022).

Recovery with the shorter arthropod NoPlant marker was less dependent on workflow than recovery with ANML or ITS. This differs from previous work on dried plant material, where shorter plant-blocking COI assays recovered more arthropod OTUs than longer assays (Krehenwinkel, Weber, Künzel, et al., 2022). In leaf litter, however, the shorter amplicon did not translate into consistently higher richness or stronger taxonomic recovery. This suggests that amplicon length alone does not determine arthropod recovery from leaf litter; primer design, plant-blocking behavior, DNA state, substrate heterogeneity, and extraction workflow likely interact. Practically, NoPlant may reduce some workflow dependence, but it should not be assumed to outperform ANML for leaf-litter arthropod biomonitoring. Moreover, it is also true that smaller amplicons contain fewer variable sites and therefore produce fewer ASVs than longer amplicons (Banerjee et al., 2026).

### 4.3 Taxonomic recovery

Higher-level taxonomic representation of arthropod orders and fungal classes also depended on workflow. For ANML, centrifugation favored several arthropod orders, including Lepidoptera, Amphipoda, Diptera, Collembola, and Hemiptera. Many of these groups plausibly contribute diffuse or surface-associated traces to litter, including tissue fragments, exuviae, frass, or fine particle-bound DNA. This supports the interpretation that high-volume wash recovery improves access to sparsely distributed animal DNA rather than simply increasing total DNA yield, consistent with broader eDNA work showing that DNA state, particle association, and sample-processing strategy influence detection (Barnes & Turner, 2016; Mauvisseau et al., 2022; Zinger et al., 2016). For ITS, the strongest taxonomic differences involved PowerSoil and included fungal classes associated with decomposing litter, spores, and hyphal biomass, including Agaricomycetes, Dothideomycetes, Sordariomycetes, Eurotiomycetes, Mucoromycetes, and Mortierellomycetes. This is consistent with direct homogenization, bead beating, and inhibitor-removal chemistry improving recovery of structurally embedded fungal DNA (Lopes et al., 2021; Schoch et al., 2012; Tedersoo et al., 2022). NoPlant showed fewer order-level differences, again illustrating that the two COI assays did not respond identically to extraction workflow. These taxonomic results matter because workflows with similar richness values may still differ in the groups they represent, a concern also noted in soil and leaf-litter metabarcoding studies where primer choice, extraction protocol, and substrate heterogeneity alter recovered taxonomic profiles (Castillo et al., 2026; Lopes et al., 2021).

### 4.4 Whole-community composition and sampling location effects

ASV-level community composition differed among extraction workflows, but sampling sites explained more variation than workflow across all markers. Previous studies have shown that soil, leaf litter, and plant-derived eDNA can recover spatial or habitat-associated variation in terrestrial communities while remaining sensitive to methodological choices. Our paired design shows both patterns in the same dataset. Most compositional variation nevertheless remained unexplained by either site or extraction workflow, consistent with the fine-scale heterogeneity of leaf-litter communities, stochastic detection in metabarcoding data, and residual technical variation.

The dispersion analyses add a qualification. Workflows differed not only in average ASV-level composition, but also in within-workflow variability. Because PERMANOVA can be influenced by both centroid shifts and dispersion differences, the multivariate results should be interpreted as evidence that workflows are not interchangeable, rather than as evidence that each workflow recovers a completely distinct community. For ITS, dispersion results were sensitive to permutation design, so those dispersion patterns should be interpreted cautiously. The broader conclusion remains that extraction workflow affected recovered composition, but sampling-site differences were the larger source of variation.

### 4.5 Workflow trade-offs and decision-making framework

The decision-space analysis and workflow framework show that extraction performance was multidimensional: no workflow maximized recovery, reproducibility, cost, safety, and scalability simultaneously **(Fig. 5**; **Fig. 6)**. This pattern is consistent with broader eDNA and metabarcoding studies showing that extraction and capture protocols can differ in DNA yield, inhibitor carryover, taxonomic recovery, reproducibility, cost, and scalability (Deiner et al., 2015; Goldberg et al., 2016; Hermans et al., 2022). In our analysis, PowerSoil emerged as the strongest default workflow for broad leaf-litter eDNA surveys because it combined high sample retention, low PCR-replicate dissimilarity, strong fungal ITS recovery, safe handling, and the shortest hands-on time (**Table 3**; **Fig. 5; Fig. S9**). Centrifugation was best supported when maximizing ANML arthropod richness, and potentially rare animal-DNA detection, was the primary goal (**Fig. 2**; **Fig. 3**; **Fig. 5**). CTAB provided the clearest cost advantage and may be useful for cost-limited scaling when phenol:chloroform handling and lower reproducibility are acceptable trade-offs (**Table 1**; **Table 3**; **Fig. 5; Fig. S9**). Filtration retained many libraries and may be useful when filter-based archiving, field-compatible processing, or compatibility with established aquatic eDNA workflows is required, but it did not exceed other methods in richness recovery, reproducibility, or cost efficiency in this analysis (**Table 3**; **Fig. 5; Fig. S9**). For multi-target surveys that require both arthropod and fungal recovery, the results support paired centrifugation and PowerSoil extractions where resources allow, or PowerSoil alone when standardized processing and reproducibility are prioritized over maximum ANML arthropod richness (**Fig. 6)**.

## 5. Limitations and next steps

Several limitations should guide the interpretation of these results. First, the study was conducted in tropical leaf litter across forest types on Oʻahu, and extraction performance may differ in systems with different litter chemistry, moisture regimes, decomposition rates, or fungal biomass. Second, extraction volume and extraction principle were partly linked by design: low-volume workflows used direct extraction from homogenized litter, whereas high-volume workflows used wash-based recovery. Therefore, we cannot fully separate the effects of input mass from the effects of recovery chemistry and handling. Third, we compared methods relative to one another but did not include an independent ground-truth community. So, while we can identify which workflow recovers more ASVs, different taxonomic profiles, or more reproducible PCR profiles, we cannot assess which method most closely represents the true community. Finally, the exposure-control results reinforce that low-biomass or open-handling eDNA workflows require explicit contamination controls, even when contaminant richness remains low.

Recommendations for future work include separating input mass from extraction principle more explicitly, for example by testing intermediate litter masses and applying direct and wash-based extraction strategies across matched inputs. Spike-in controls or mock communities have also proven useful to quantify absolute DNA recovery and inhibition, although realistic mock communities for leaf litter will be difficult to design. Finally, testing the same workflows across temperate, tropical, dry, and wet litter systems would determine whether the marker-specific patterns observed here are general or habitat-dependent.

## 6. Conclusion

All leaf-litter eDNA workflows tested here recovered site-level differences among forest-floor communities, but biodiversity recovery depended strongly on both extraction workflow and marker (taxon target). Our results suggest using high-volume workflow (e.g., Centrifugation) can maximize arthropod detection, and low-volume workflow (e.g., PowerSoil) can maximize the fungal detection. CTAB was cost-effective but limited by handling hazardous materials. Filtration was not favored by measured recovery or cost efficiency, although it may remain useful when filter-based storage or protocol compatibility is required. The results can be interpreted in terms of fraction recovery and lead to a well-supported decision tree for choice of method (**Fig. 6**). Standardizing or explicitly calibrating extraction workflows will be essential for comparing leaf-litter eDNA data among taxa, sites, studies, and monitoring programs.

### Author Contributions

Sven Weber (SWE), Pritam Banerjee (PBA), Edoardo Scali (ESC), Alex Augustus Farrow (AAF), Alex Maria Boren (AMB), Walter Thomas Russell III (WTR), Rosemary Gillespie (RGG), Natalie R. Graham (NRG) and George Roderick (GKR) contributed to the study. SWE, PBA, AAF, NRG and GKR conceived the ideas and designed the methodology. SWE, AAF, WTR, RGG, NRG and GKR collected the samples. SWE, PBA, AAF, AMB and contributed to laboratory work. SWE and ESC conducted the bioinformatic processing. SWE and ESC analysed the data. SWE led the writing of the manuscript, with substantial contributions from RGG, GKR and NRG. All authors contributed critically to the drafts and gave final approval for publication, SWE and PBA are shared first authors.

### Data Availability

Raw sequences, processed ASV tables, sample metadata, and analysis scripts will be available from Dryad at: https://datadryad.org/dataset/doi:10.5061/dryad.7sqv9s581

The full data citation will be included in the reference list upon acceptance.After acceptance code will be reachable at gith: https://github.com/sven9r/SERDP_leaflitter_method

## References

Abarenkov, K., Nilsson, R. H., Larsson, K.-H., Taylor, A. F., May, T. W., Frøslev, T. G., Pawlowska, J., Lindahl, B., Põldmaa, K., & Truong, C. (2024). The UNITE database for molecular identification and taxonomic communication of fungi and other eukaryotes: Sequences, taxa and classifications reconsidered. Nucleic Acids Research, 52(D1), D791–D797.

Anderson, J. M. (1975). Succession, Diversity and Trophic Relationships of Some Soil Animals in Decomposing Leaf Litter. The Journal of Animal Ecology, 44(2), 475. 10.2307/3607

Banerjee, P., Al-Bayer, S., Calaor, J., Weber, S., Graham, N. R., Andersen, J. C., Economo, E. P., Kennedy, S., Krehenwinkel, H., Gillespie, R. G., Roderick, G. K., Rogers, H. S., & Puliafico, K. P. (2026). Comparison of Environmental DNA and Bulk DNA Metabarcoding for Assessing Terrestrial Arthropod Diversity Across Three Habitat Types on Guam. Molecular Ecology Resources, 26(5), e70172. 10.1111/1755-0998.70172

Banerjee, P., Maity, J. P., Chatterjee, N., Weber, S., Dey, G., Sharma, R. K., & Chen, C. (2025). Harnessing Environmental DNA to Explore Frugivorous Interactions: A Case Study in Papaya (Carica papaya) and Pineapple (Ananas comosus). Environmental DNA, 7(5), e70196.

Barnes, M. A., & Turner, C. R. (2016). The ecology of environmental DNA and implications for conservation genetics. Conservation Genetics, 17(1), 1–17. 10.1007/s10592-015-0775-4

Basset, Y., Donoso, D. A., Hajibabaei, M., Wright, M. T. G., Perez, K. H. J., Lamarre, G. P. A., De León, L. F., Palacios-Vargas, J. G., Castaño-Meneses, G., Rivera, M., Perez, F., Bobadilla, R., Lopez, Y., Ramirez, J. A., & Barrios, H. (2020). Methodological considerations for monitoring soil/litter arthropods in tropical rainforests using DNA metabarcoding, with a special emphasis on ants, springtails and termites. Metabarcoding and Metagenomics, 4, e58572. 10.3897/mbmg.4.58572

Basset, Y., Hajibabaei, M., Wright, M. T. G., Castillo, A. M., Donoso, D. A., Segar, S. T., Souto-Vilarós, D., Soliman, D. Y., Roslin, T., Smith, M. A., Lamarre, G. P. A., De León, L. F., Decaëns, T., Palacios-Vargas, J. G., Castaño-Meneses, G., Scheffrahn, R. H., Rivera, M., Perez, F., Bobadilla, R., … Barrios, H. (2022). Comparison of traditional and DNA metabarcoding samples for monitoring tropical soil arthropods (Formicidae, Collembola and Isoptera). Scientific Reports, 12(1), 10762. 10.1038/s41598-022-14915-2

Bates, D., Mächler, M., Bolker, B., & Walker, S. (2015). Fitting Linear Mixed-Effects Models Using **lme4**. Journal of Statistical Software, 67(1). 10.18637/jss.v067.i01

Benjamini, Y., & Hochberg, Y. (1995). Controlling the False Discovery Rate: A Practical and Powerful Approach to Multiple Testing. Journal of the Royal Statistical Society Series B: Statistical Methodology, 57(1), 289–300. 10.1111/j.2517-6161.1995.tb02031.x

Buchner, D., Haase, P., & Leese, F. (2021). Wet grinding of invertebrate bulk samples–a scalable and cost-efficient protocol for metabarcoding and metagenomics. Metabarcoding and Metagenomics, 5, e67533. 10.3897/mbmg.5.67533

Camacho, C., Coulouris, G., Avagyan, V., Ma, N., Papadopoulos, J., Bealer, K., & Madden, T. L. (2009). BLAST+: Architecture and applications. BMC Bioinformatics, 10(1), 421. 10.1186/1471-2105-10-421

Castillo, A. H., Jacobs, S., Steinke, D., & Smith, M. A. (2026). Assessment of Leaf-Litter Invertebrate Biodiversity Using High Throughput Sequencing. bioRxiv, 2026–04.

Deagle, B. E., Jarman, S. N., Coissac, E., Pompanon, F., & Taberlet, P. (2014). DNA metabarcoding and the cytochrome *c* oxidase subunit I marker: Not a perfect match. Biology Letters, 10(9), 20140562. 10.1098/rsbl.2014.0562

Deiner, K., Walser, J.-C., Mächler, E., & Altermatt, F. (2015). Choice of capture and extraction methods affect detection of freshwater biodiversity from environmental DNA. Biological Conservation, 183, 53–63. 10.1016/j.biocon.2014.11.018

Doyle, J. J., & Doyle, J. L. (1987). A rapid DNA isolation procedure for small quantities of fresh leaf tissue. Phytochemical Bulletin.

Elbrecht, V., & Leese, F. (2017). Validation and development of COI metabarcoding primers for freshwater macroinvertebrate bioassessment. Frontiers in Environmental Science, 5(APR), 11–11. 10.3389/fenvs.2017.00011

Folmer, O., Hoeh, W., Black, M., & Vrijenhoek, R. (1994). Conserved primers for PCR amplification of mitochondrial DNA from different invertebrate phyla. Molecular Marine Biology and Biotechnology, 3(5), 294–299.

Gardes, M., & Bruns, T. D. (1993). ITS primers with enhanced specificity for basidiomycetes - application to the identification of mycorrhizae and rusts. Molecular Ecology, 2(2), 113–118. 10.1111/j.1365-294X.1993.tb00005.x

Goldberg, C. S., Turner, C. R., Deiner, K., Klymus, K. E., Thomsen, P. F., Murphy, M. A., Spear, S. F., McKee, A., Oyler-McCance, S. J., Cornman, R. S., Laramie, M. B., Mahon, A. R., Lance, R. F., Pilliod, D. S., Strickler, K. M., Waits, L. P., Fremier, A. K., Takahara, T., Herder, J. E., & Taberlet, P. (2016). Critical considerations for the application of environmental DNA methods to detect aquatic species. Methods in Ecology and Evolution, 7(11), 1299–1307. 10.1111/2041-210X.12595

Hermans, S. M., Lear, G., Buckley, T. R., & Buckley, H. L. (2022). Environmental DNA sampling detects between-habitat variation in soil arthropod communities, but is a poor indicator of fine-scale spatial and seasonal variation. Ecological Indicators, 140, 109040. 10.1016/j.ecolind.2022.109040

Horton, D. J., Kershner, M. W., & Blackwood, C. B. (2017). Suitability of PCR primers for characterizing invertebrate communities from soil and leaf litter targeting metazoan 18S ribosomal or cytochrome oxidase I (COI) genes. European Journal of Soil Biology, 80, 43–48. 10.1016/j.ejsobi.2017.04.003

Jusino, M. A., Banik, M. T., Palmer, J. M., Wray, A. K., Xiao, L., Pelton, E., Barber, J. R., Kawahara, A. Y., Gratton, C., & Peery, M. Z. (2019). An improved method for utilizing high-throughput amplicon sequencing to determine the diets of insectivorous animals. Molecular Ecology Resources, 19(1), 176–190. 10.1111/1755-0998.12951

Krehenwinkel, H., Weber, S., Broekmann, R., Melcher, A., Hans, J., Wolf, R., Hochkirch, A., Kennedy, S. R., Koschorreck, J., & Künzel, S. (2022). Environmental DNA from archived leaves reveals widespread temporal turnover and biotic homogenization in forest arthropod communities. Elife, 11, e78521.

Krehenwinkel, H., Weber, S., Künzel, S., & Kennedy, S. R. (2022). The bug in a teacup—Monitoring arthropod–plant associations with environmental DNA from dried plant material. Biology Letters, 18(6), 20220091. 10.1098/rsbl.2022.0091

Lange, V., Böhme, I., Hofmann, J., Lang, K., Sauter, J., Schöne, B., Paul, P., Albrecht, V., Andreas, J. M., Baier, D. M., Nething, J., Ehninger, U., Schwarzelt, C., Pingel, J., Ehninger, G., & Schmidt, A. H. (2014). Cost-efficient high-throughput HLA typing by MiSeq amplicon sequencing. BMC Genomics, 15(1), 63. 10.1186/1471-2164-15-63

Longmire, J. L., Maltbie, M., & Baker, R. J. (1997). Use of" lysis buffer" in DNA isolation and its implication for museum collections.

Lopes, C. M., Baêta, D., Sasso, T., Vanzetti, A., Raquel Zamudio, K., Taberlet, P., & Haddad, C. F. B. (2021). Power and limitations of environmental DNA metabarcoding for surveying leaf litter eukaryotic communities. Environmental DNA, 3(3), 528–540. 10.1002/edn3.142

Mauvisseau, Q., Harper, L. R., Sander, M., Hanner, R. H., Kleyer, H., & Deiner, K. (2022). The Multiple States of Environmental DNA and What Is Known about Their Persistence in Aquatic Environments. Environmental Science & Technology, 56(9), 5322–5333. 10.1021/acs.est.1c07638

McKee, A. M., Spear, S. F., & Pierson, T. W. (2015). The effect of dilution and the use of a post-extraction nucleic acid purification column on the accuracy, precision, and inhibition of environmental DNA samples. Biological Conservation, 183, 70–76.

Oksanen, J., Blanchet, F. G., Kindt, R., Legendre, P., Minchin, P. R., O’hara, R., Simpson, G. L., Solymos, P., Stevens, M. H. H., & Wagner, H. (2013). Package ‘vegan.’ *Community Ecology Package*, Version, 2(9), 1–295.

Osono, T. (2007). Ecology of ligninolytic fungi associated with leaf litter decomposition. Ecological Research, 22(6), 955–974. 10.1007/s11284-007-0390-z

Prescott, C. E., & Vesterdal, L. (2021). Decomposition and transformations along the continuum from litter to soil organic matter in forest soils. Forest Ecology and Management, 498, 119522. 10.1016/j.foreco.2021.119522

R Core Team. (2024). R: A Language and Environment for Statistical Computing [Computer software]. R Foundation for Statistical Computing. https://www.R-project.org/

Rognes, T., Flouri, T., Nichols, B., Quince, C., & Mahé, F. (2016). VSEARCH: a versatile open source tool for metagenomics. PeerJ, 4, e2584. 10.7717/peerj.2584

Ruppert, L.-S., Staab, M., Klingenfuß, S., Rappa, N. J., Frey, J., & Segelbacher, G. (2023). Leaf litter arthropods show little response to structural retention in a Central European forest. Biodiversity and Conservation, 32(12), 3973–3990. 10.1007/s10531-023-02677-w

Salter, S. J., Cox, M. J., Turek, E. M., Calus, S. T., Cookson, W. O., Moffatt, M. F., Turner, P., Parkhill, J., Loman, N. J., & Walker, A. W. (2014). Reagent and laboratory contamination can critically impact sequence-based microbiome analyses. BMC Biology, 12(1), 87. 10.1186/s12915-014-0087-z

Schloss, P. D. (2024). Waste not, want not: Revisiting the analysis that called into question the practice of rarefaction. Msphere, 9(1), e00355–23.

Schoch, C. L., Seifert, K. A., Huhndorf, S., Robert, V., Spouge, J. L., Levesque, C. A., Chen, W., Fungal Barcoding Consortium, Fungal Barcoding Consortium Author List, Bolchacova, E., Voigt, K., Crous, P. W., Miller, A. N., Wingfield, M. J., Aime, M. C., An, K.-D., Bai, F.-Y., Barreto, R. W., Begerow, D., … Schindel, D. (2012). Nuclear ribosomal internal transcribed spacer (ITS) region as a universal DNA barcode marker for *Fungi*. Proceedings of the National Academy of Sciences, 109(16), 6241–6246. 10.1073/pnas.1117018109

Schöneberg, Y. (2023). Yschoeneberg/blast2taxonomy: V1. 3.4.

Sidstedt, M., Rådström, P., & Hedman, J. (2020). PCR inhibition in qPCR, dPCR and MPS—mechanisms and solutions. Analytical and Bioanalytical Chemistry, 412(9), 2009.

Stothut, M., Mahla, L., Backes, L., Weber, S., Avazzadeh, A., Moradmand, M., & Krehenwinkel, H. (2024). Recovering plant-associated arthropod communities by eDNA metabarcoding historical herbarium specimens. Current Biology, 34(18), 4318–4324. 10.1016/j.cub.2024.07.100

Tedersoo, L., Bahram, M., Zinger, L., Nilsson, R. H., Kennedy, P. G., Yang, T., Anslan, S., & Mikryukov, V. (2022). Best practices in metabarcoding of fungi: From experimental design to results. Molecular Ecology, 31(10), 2769–2795. 10.1111/mec.16460

Weber, S., Hutchins, L., Banerjee, P., Callaghan, W., Farrow, A. A., Andersen, J., Gillespie, R., & Roderick, G. (2026). Topography structures of arthropod communities revealed by leaf-derived environmental DNA on O’ahu, Hawai’i. bioRxiv, 2026–04.

Weber, S., Stothut, M., Mahla, L., Kripp, A., Hirschler, L., Lenz, N., Junker, A., Künzel, S., & Krehenwinkel, H. (2024). Plant-derived environmental DNA complements diversity estimates from traditional arthropod monitoring methods but outperforms them detecting plant–arthropod interactions. Molecular Ecology Resources, 24(2), e13900. 10.1111/1755-0998.13900

White, T. J., Bruns, T., Lee, S., & Taylor, J. (1990). Amplification and direct sequencing of fungal ribosomal RNA genes for phylogenetics. PCR Protocols: A Guide to Methods and Applications, 18(1), 315–322.

Yang, C., Wang, X., Miller, J. A., De Blécourt, M., Ji, Y., Yang, C., Harrison, R. D., & Yu, D. W. (2014). Using metabarcoding to ask if easily collected soil and leaf-litter samples can be used as a general biodiversity indicator. Ecological Indicators, 46, 379–389. 10.1016/j.ecolind.2014.06.028

Young, M. R., & Hebert, P. D. N. (2022). Unearthing soil arthropod diversity through DNA metabarcoding. PeerJ, 10, e12845. 10.7717/peerj.12845

Zhang, J., Kobert, K., Flouri, T., & Stamatakis, A. (2014). PEAR: a fast and accurate Illumina Paired-End reAd mergeR. Bioinformatics, 30(5), 614–620. 10.1093/bioinformatics/btt593

Zinger, L., Chave, J., Coissac, E., Iribar, A., Louisanna, E., Manzi, S., Schilling, V., Schimann, H., Sommeria-Klein, G., & Taberlet, P. (2016). Extracellular DNA extraction is a fast, cheap and reliable alternative for multi-taxa surveys based on soil DNA. Soil Biology and Biochemistry, 96, 16–19. 10.1016/j.soilbio.2016.01.008

