## Supplemental for "Extraction workflow determines marker-specific recovery and reproducibility in leaf-litter eDNA metabarcoding"

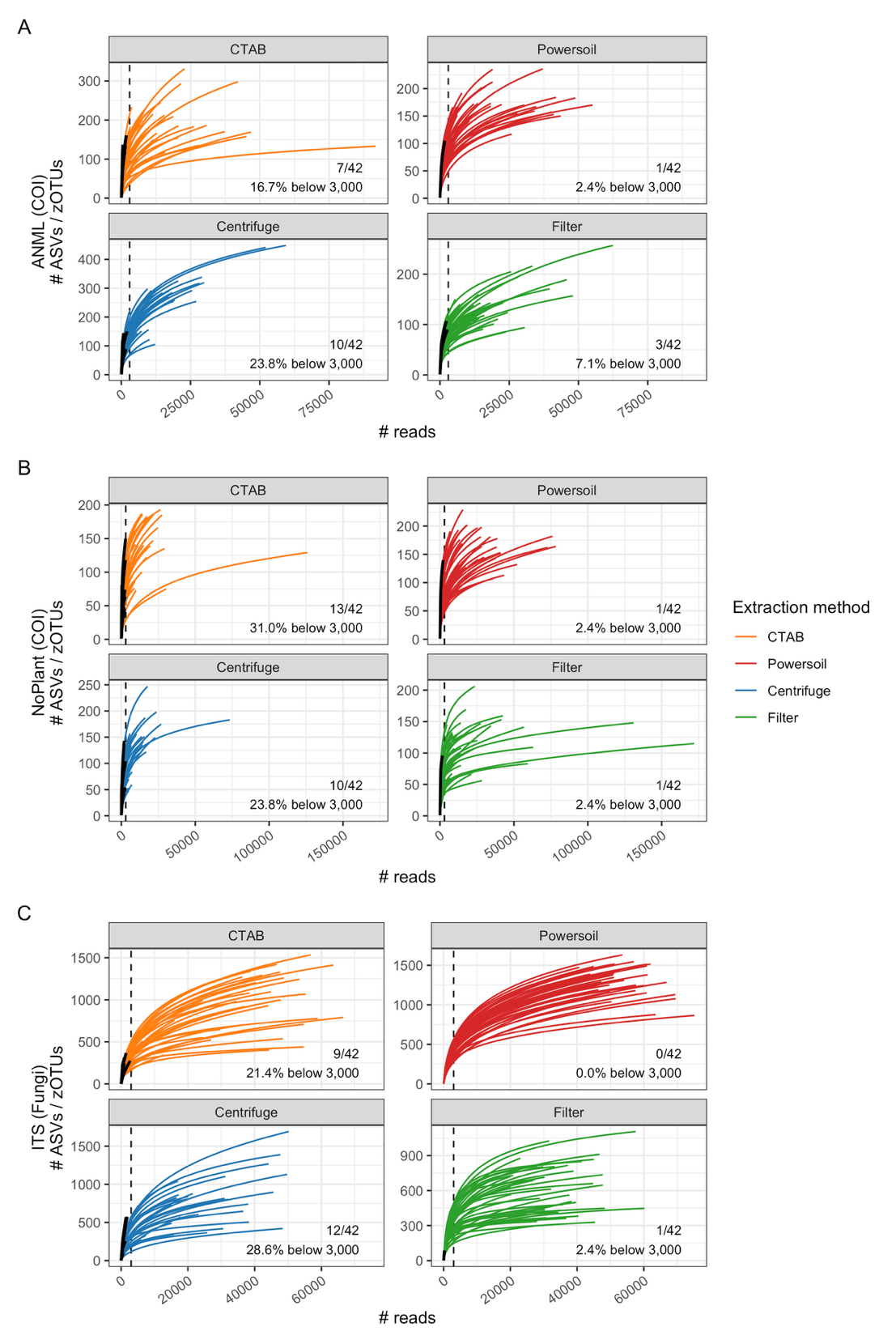

**Figure S1.** Rarefaction curves showing the accumulation of ASV richness with sequencing depth across extraction methods and markers at the biological unit level. Panels represent (A) ANML (COI), (B) NoPlant (COI), and (C) ITS (fungi). Each line corresponds to a biological unit, colored by extraction method (Centrifugation, CTAB, Filter, PowerSoil). The dashed vertical line indicates the rarefaction threshold (3,000 reads). Numbers within panels indicate the number and proportion of samples falling below this threshold.

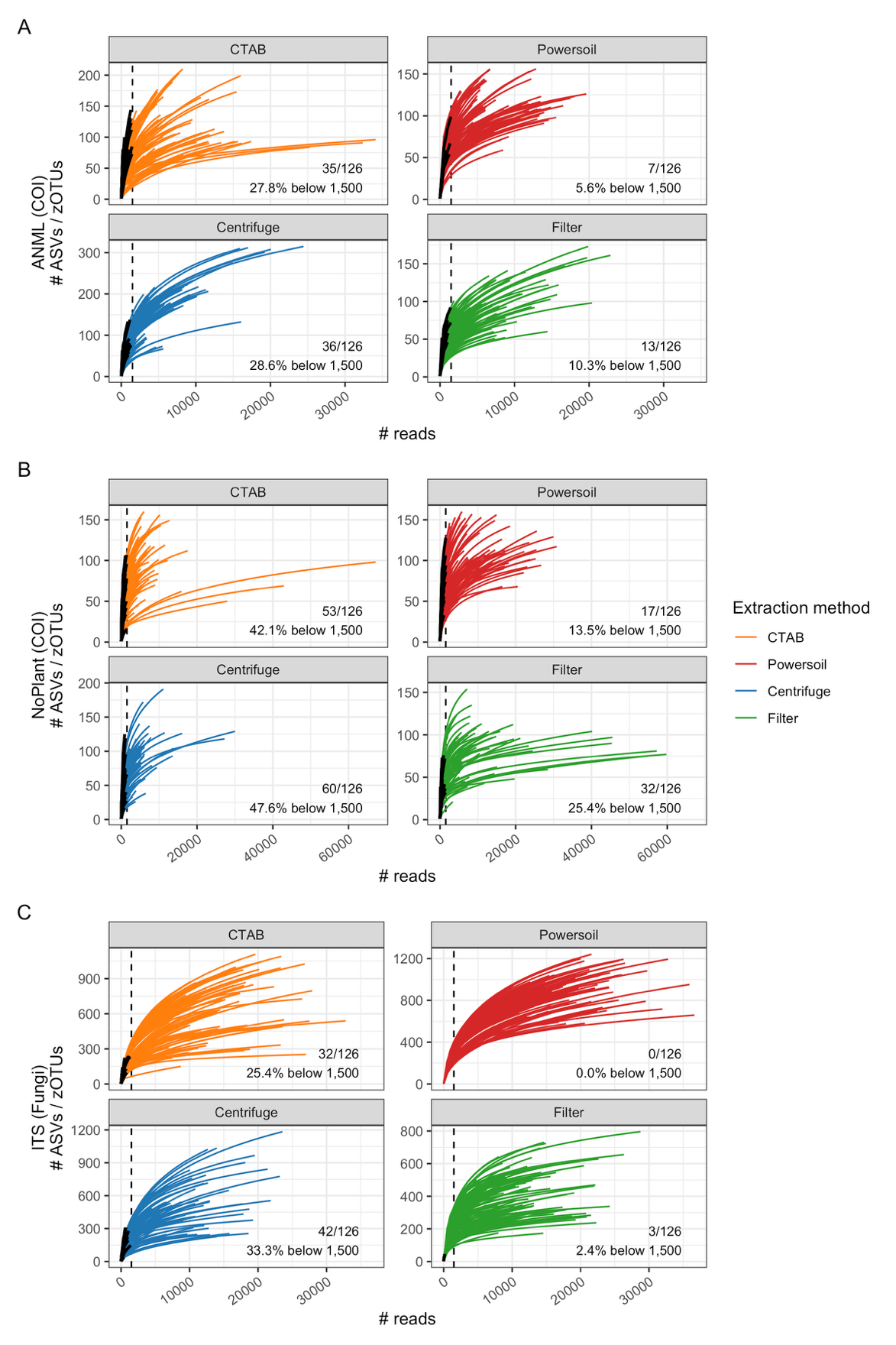

**Figure S2.** Rarefaction curves showing ASV richness as a function of sequencing depth across extraction methods and markers at the PCR replicate level. Panels represent (A) ANML (COI), (B) NoPlant (COI), and (C) ITS (fungi). Each line corresponds to a PCR replicate, colored by extraction method. The dashed vertical line indicates the rarefaction threshold (1,500 reads). Numbers within panels indicate the number and proportion of replicates below this threshold.
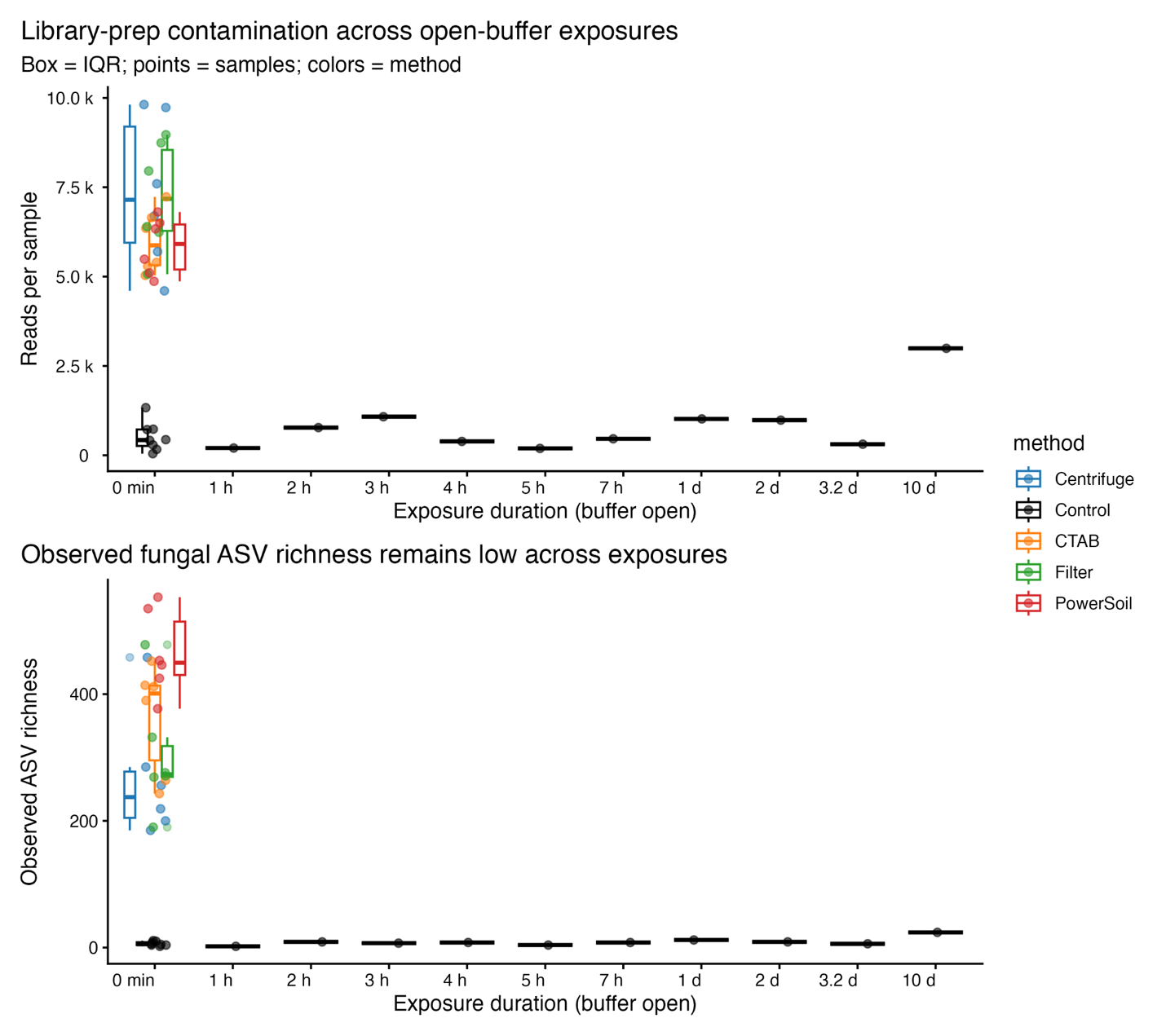
**Figure S3.** Library preparation contamination and its effect on sequencing output and observed fungal richness across increasing exposure times of open extraction buffers. (Top) Total reads per sample across exposure durations for each extraction method and controls. (Bottom) Observed fungal ASV richness across the same exposure gradient. Boxes represent interquartile ranges, with points indicating individual samples.
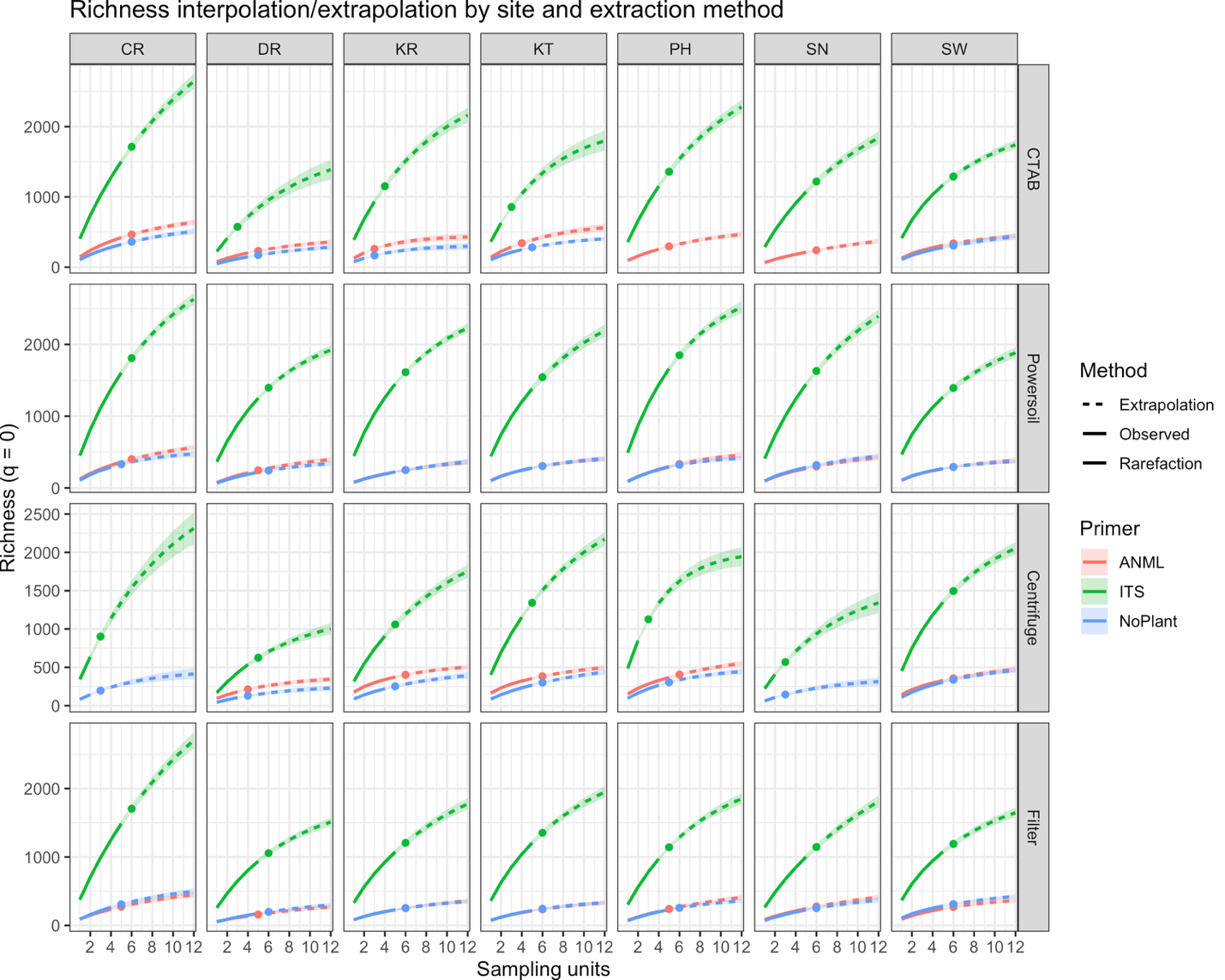

**Figure S4.** Sample-size-based rarefaction and extrapolation curves (Hill number q = 0) for ASV richness across sites and extraction methods. Rows correspond to extraction methods and columns to sampling sites. Colors indicate primer sets (ANML, ITS, NoPlant). Solid lines represent interpolation (rarefaction), dashed lines represent extrapolation, and points indicate observed richness. Shaded areas denote 95% confidence intervals.

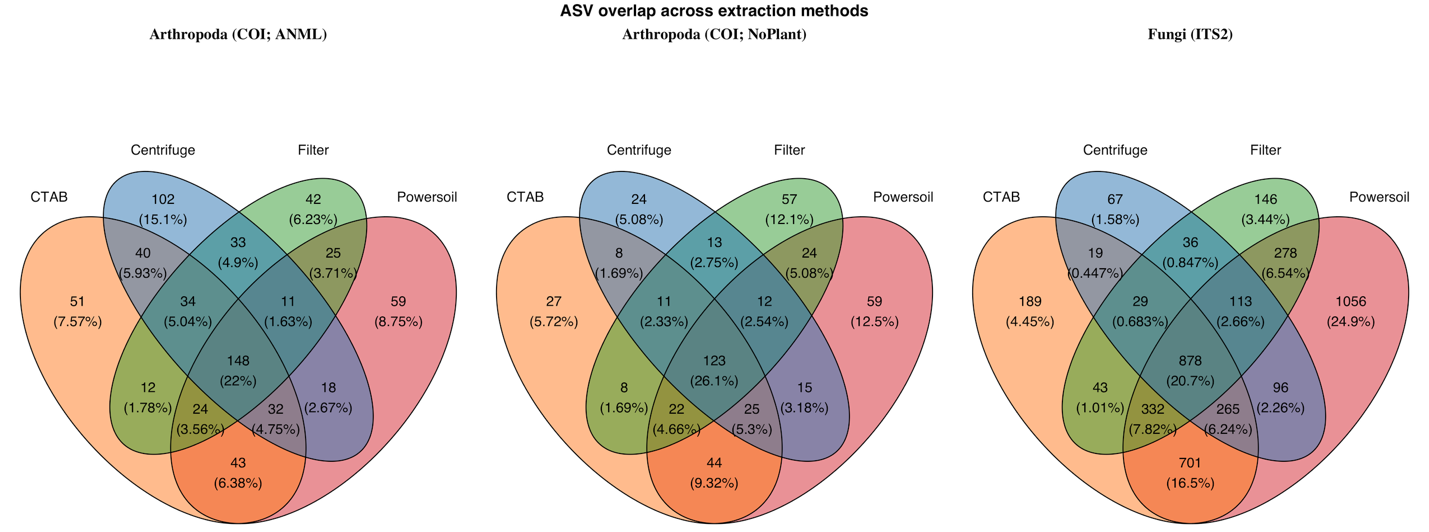

**Figure S5. ASV overlap across extraction methods for each marker**
Venn diagrams show overlap in recovered ASVs among the four extraction workflows (CTAB, PowerSoil, centrifugation, and filtration) using three different primers (ANML COI arthropods, NoPlant COI arthropods, and ITS fungi). Numbers indicate ASVs assigned to each intersection, with percentages showing the proportion of the total ASV pool represented by that region.

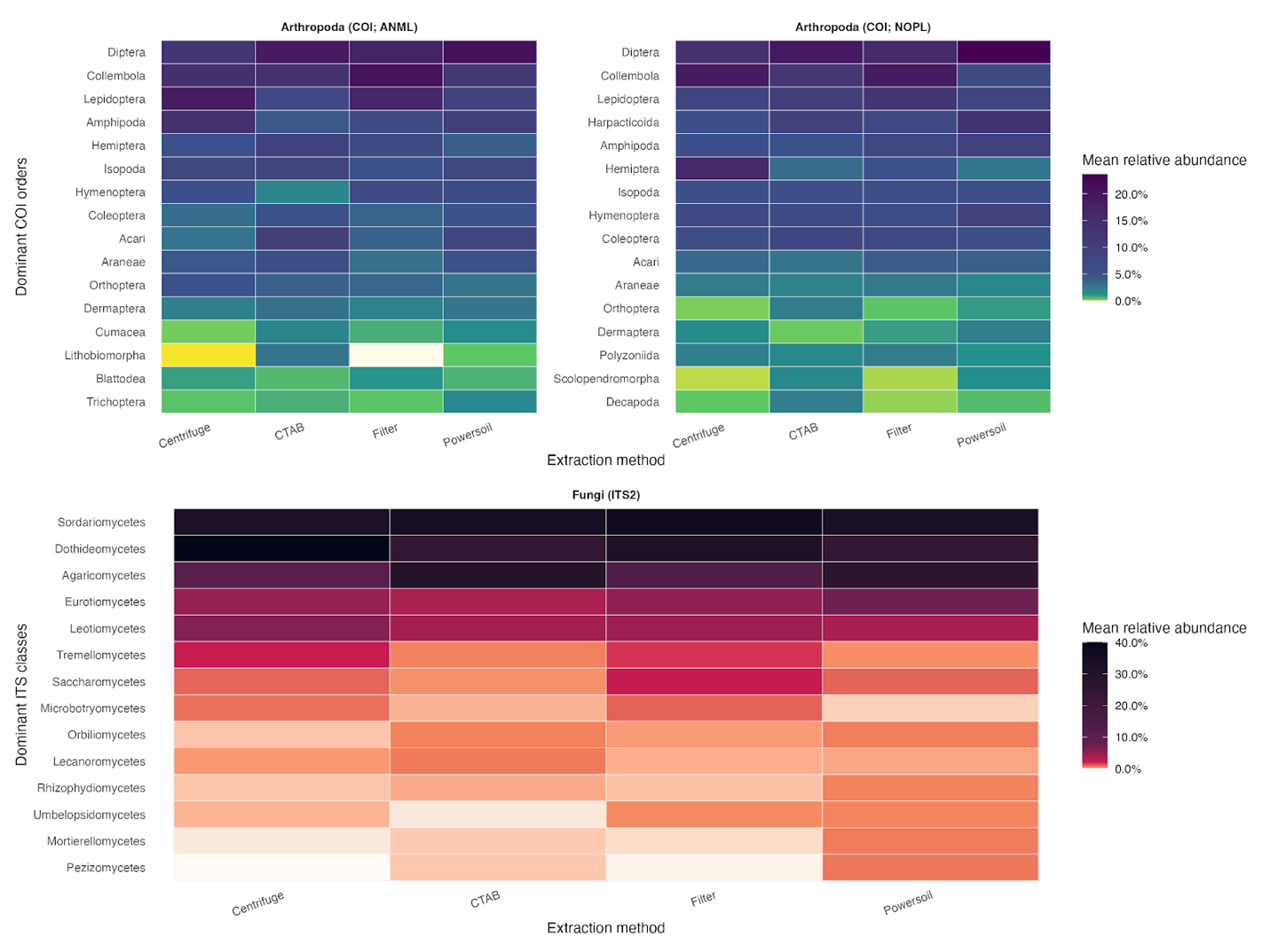

**Figure S6.** Mean relative abundance of dominant taxonomic groups across extraction methods and markers. Top panels show dominant arthropod orders for ANML and NoPlant markers; bottom panel shows dominant fungal classes for ITS. Colors indicate mean relative abundance across samples for each extraction method.

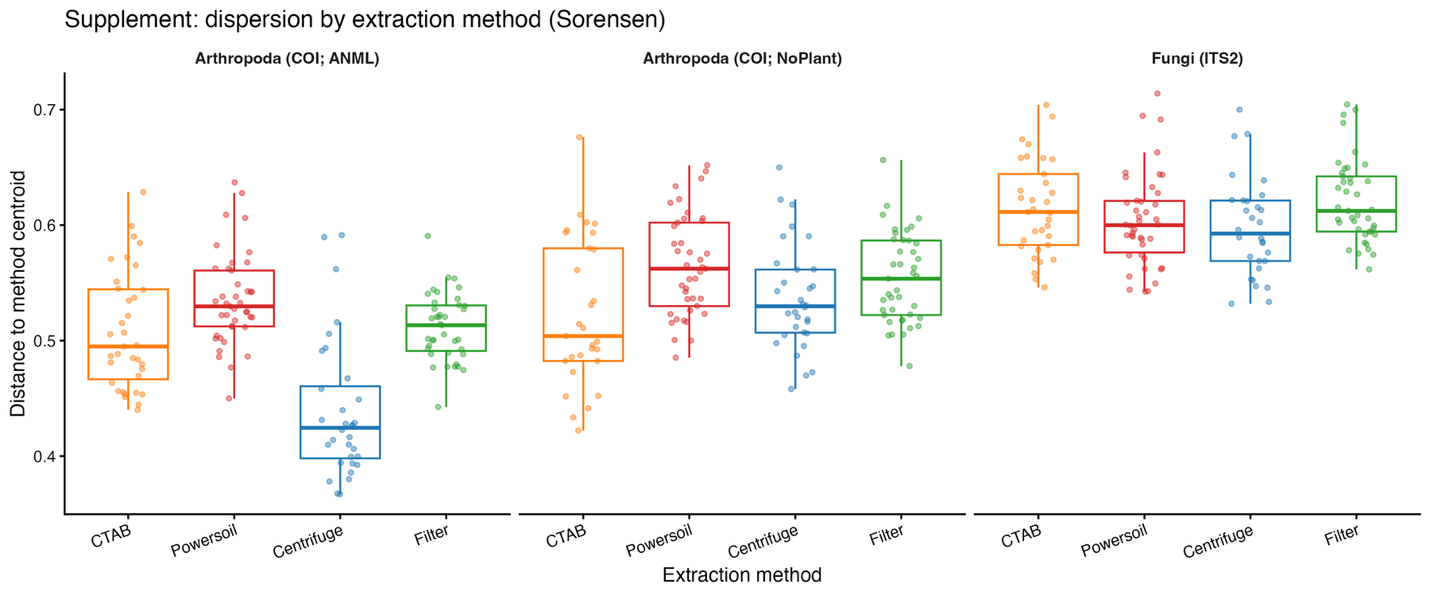

**Figure S7.** Within-method dispersion of community composition measured as distance to centroid based on Sørensen dissimilarity. Panels represent (left to right) ANML (COI), NoPlant (COI), and ITS (fungi). Points correspond to biological units, colored by extraction method. Boxes indicate interquartile ranges with medians. Higher values indicate greater dispersion among samples within an extraction method.

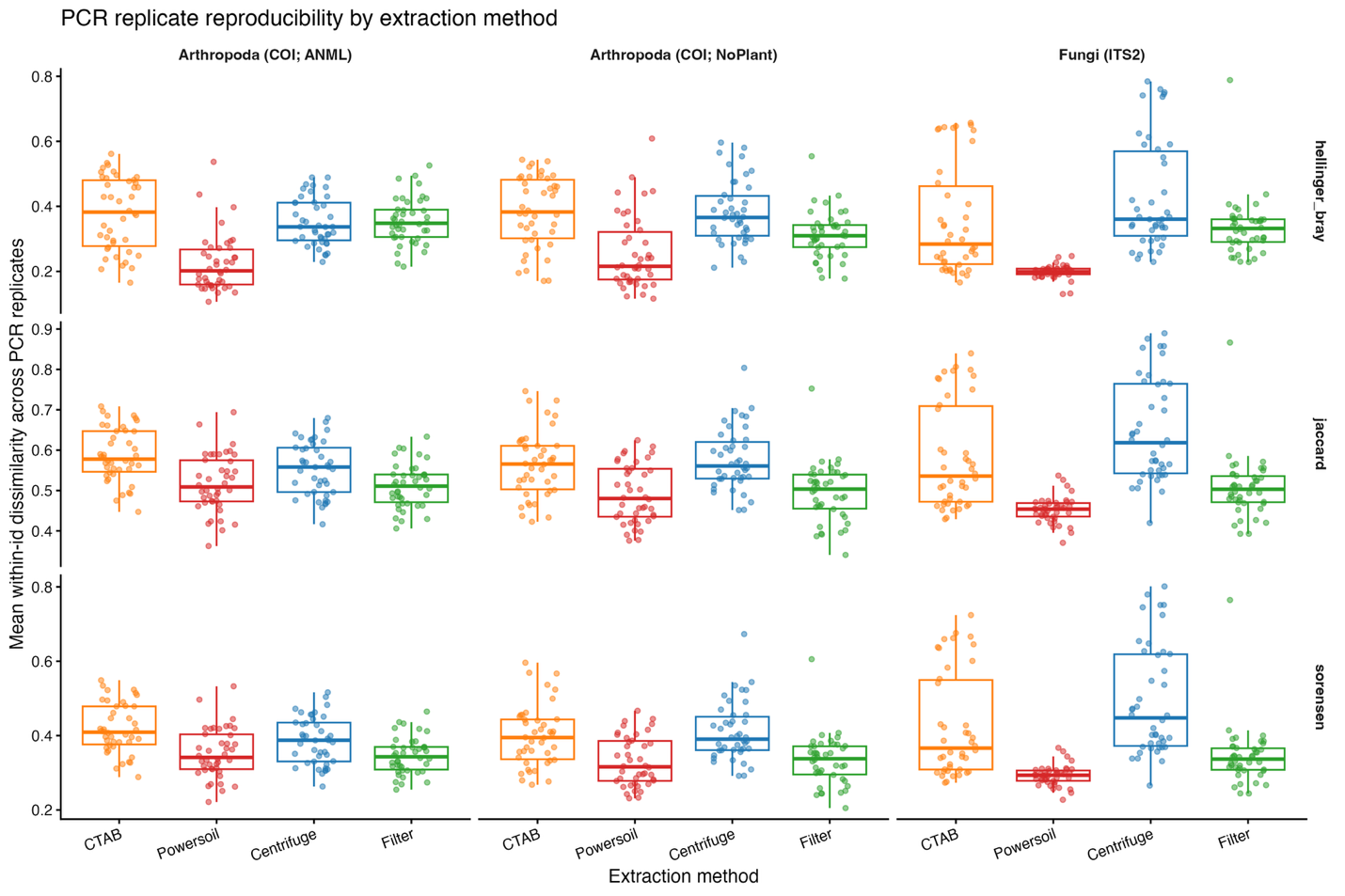

**Figure S8**. Within-sample reproducibility across PCR replicates measured as mean pairwise dissimilarity for each extraction method. Panels represent markers (columns: ANML, NoPlant, ITS) and distance metrics (rows: Hellinger–Bray, Jaccard, Sørensen). Points represent biological units; boxplots summarize distributions. Lower dissimilarity indicates higher reproducibility among PCR replicates.

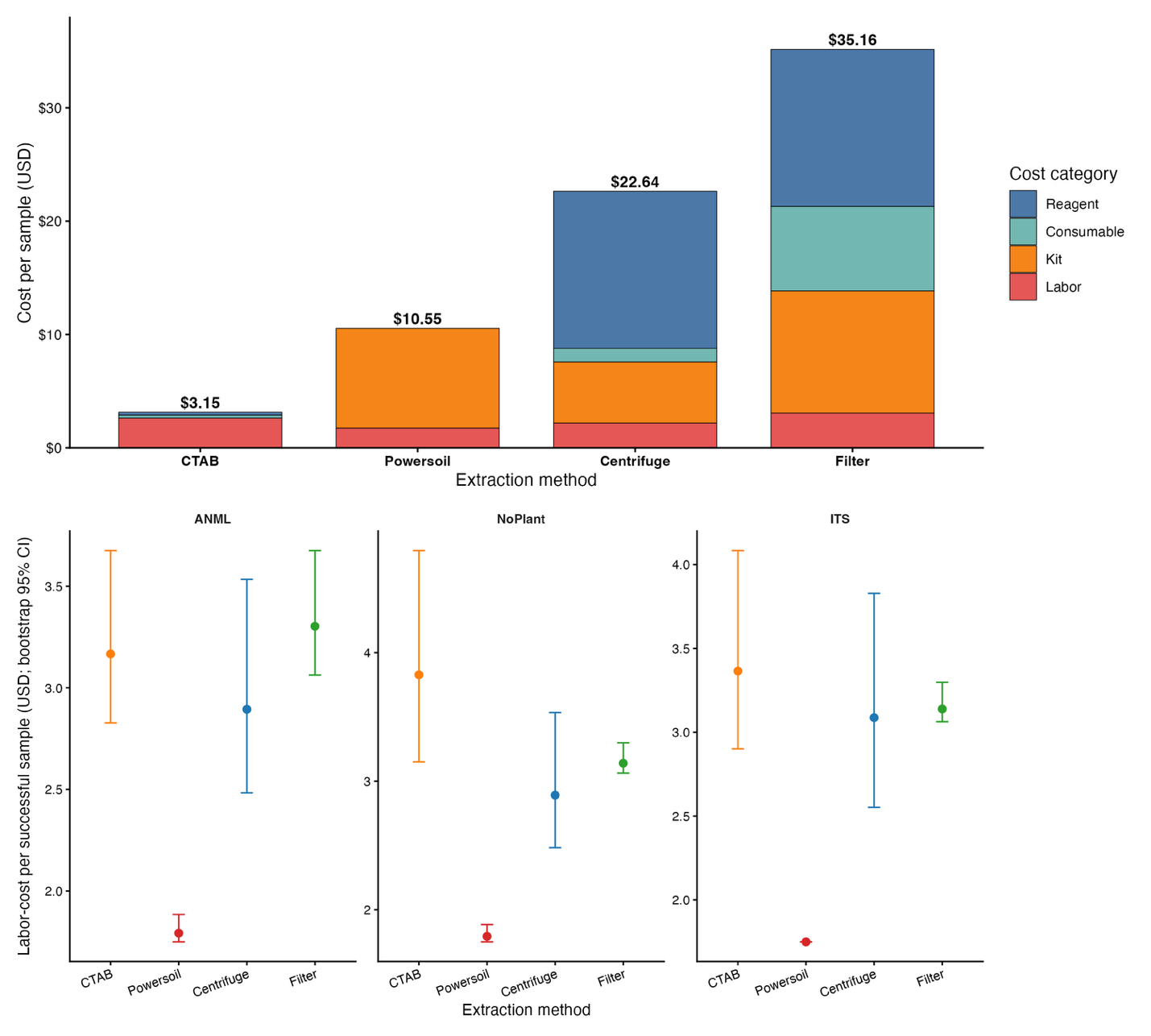

**Figure S9. Cost structure and total processing cost of the four extraction methods.** The upper panel shows per-sample cost partitioned into labor, kit, consumable, and reagent components. Lower panels summarize total cost per sample and per batch. CTAB had the lowest estimated cost ($3.15 per sample; $75.68 per batch), followed by PowerSoil ($10.55 per sample; $253.20 per batch), Centrifugation ($22.64 per sample; $543.43 per batch), and Filter ($35.16 per sample; $843.80 per batch). Reagent costs dominated the Centrifugation and Filter workflows, whereas kit costs dominated PowerSoil. These estimates provide a practical context for interpreting tradeoffs between extraction performance and scalability.

Table S1:

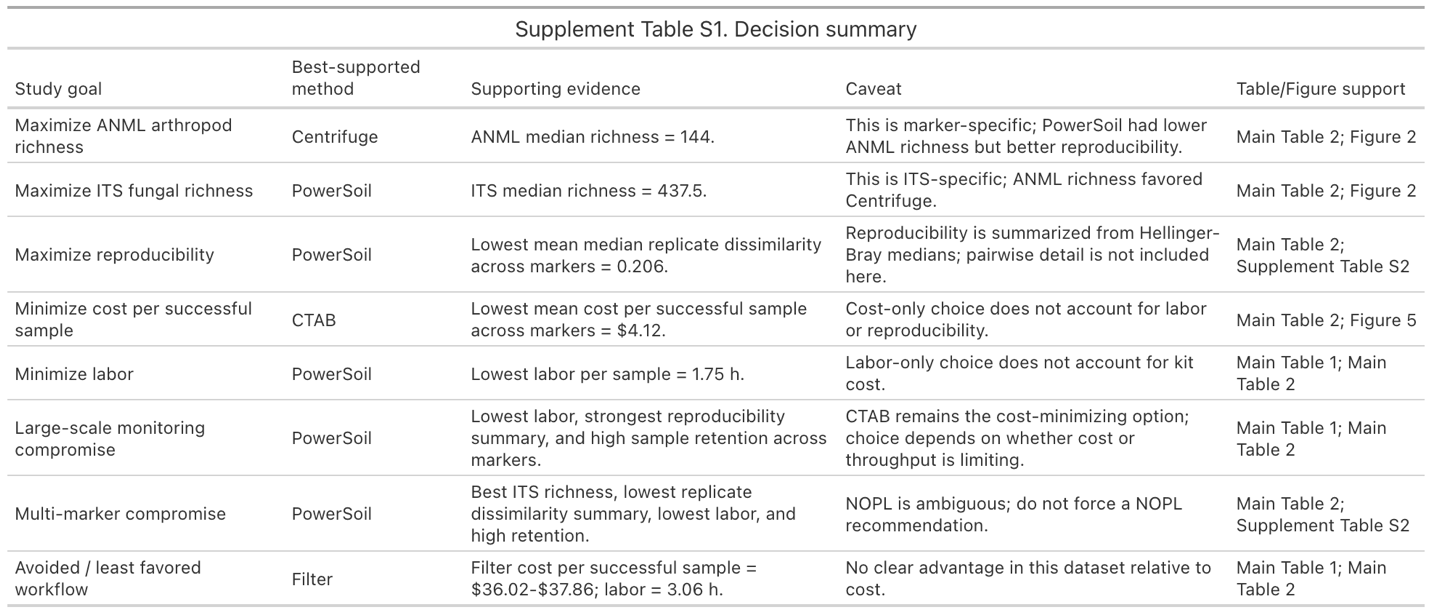

Table S2

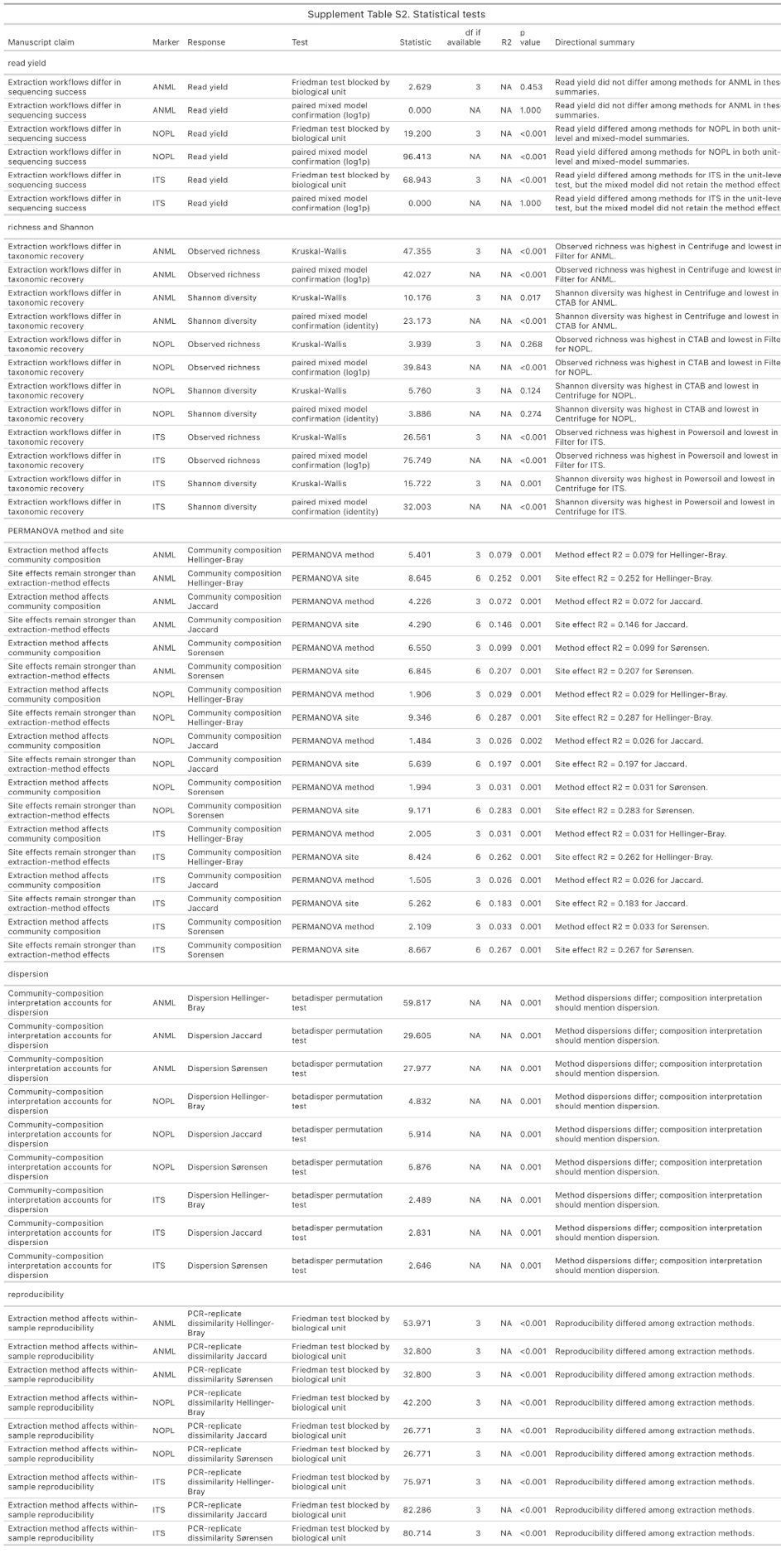
